# Spatial Organellomics Maps Cell State Diversity and Metabolic Adaptation in Tissues

**DOI:** 10.1101/2024.12.06.627285

**Authors:** Raghabendra Adhikari, Alexander Hillsley, Alana Dowdell Johnson, Shihong Max Gao, Isabel Espinosa-Medina, Jan Funke, Daniel Feliciano

**Affiliations:** Janelia Research Campus, Howard Hughes Medical Institute; Ashburn, VA, 20147; USA; Chan Zuckerberg Biohub, San Francisco, CA, USA; Janelia-Meyerhoff Undergraduate Scholars Program 2024

**Author notes:** These authors contributed equally to this work (Raghabendra Adhikari, Alexander Hillsley).

## Abstract

Cell state diversity drives tissue adaptability, repair, and disease resilience, but fully capturing this cellular complexity remains a challenge. Most current approaches rely on transcriptional profiling and often overlook functional insights embedded in organelle structure, key indicators of cellular metabolism and stress. Here, we introduce *spatial Organellomics* (*sOrganellomics*), an imaging workflow integrating automated segmentation with machine learning to classify and spatially map cell states based on multi-organelle features. Across metabolically specialized organs, these features distinguish organ-specific cellular identities. In the liver, *sOrganellomics* reveals a previously unrecognized hepatocyte organization: different hepatocyte states form intermixed communities rather than classical homogeneous graded zones, a pattern disrupted by nutritional stress. Intravital imaging further links fasting-associated architectural remodeling to altered mitochondrial membrane potential and bioenergetic heterogeneity *in vivo*. This approach maps metabolic transitions and early disease progression, establishing organelle architecture as a scalable readout of functional cell state diversity in tissues.

## Introduction

Organs are composed of a dynamic mosaic of cells in diverse functional states, each integrating cues from its local microenvironment. Defining this heterogeneity is essential for understanding how tissues coordinate physiology, adapt to stress, and lose resilience in disease (*1–3*). Over the past decade, large-scale efforts to map cellular diversity have relied primarily on transcriptional profiling, revealing extensive variation in gene expression programs across tissues (*4–10*). Yet gene expression captures only one layer of cellular state. Many functional properties of cells are executed through the organization of intracellular compartments, known as organelles, which integrate transcriptional, translational, and post-translational regulation in space and time. Organelle architecture therefore represents an important dimension of cell-state biology (*11–16*). However, despite its potential to reveal functional cellular heterogeneity, technical challenges in imaging and analyzing multiple organelles *in situ* have limited the use of organelle architecture as a scalable readout of cellular states.

Organelle architecture encompasses the abundance, morphology, and spatial distribution of intracellular compartments. These properties are dynamic and remodel in response to metabolic demands and cellular stress (*11–31*). For example, mitochondrial networks elongate during nutrient scarcity to enhance bioenergetic efficiency, whereas peroxisomes and lipid droplets alter their abundance and distribution during redox and metabolic stress (*17–27*). In the liver remodeling of endoplasmic reticulum (ER) architecture during obesity disrupts protein secretion and impairs systemic glucose homeostasis, effects that can be reversed by restoring ER structural organization (*18*). Such findings indicate that organelle architecture is closely coupled to cellular function and can influence it directly. Quantitative analysis of organelle organization could therefore provide a functional readout of cell state in tissues that complements molecular profiling.

Recent advances in multiplexed fluorescence microscopy and machine-learning-based image analysis have begun to make this possible in cultured cells. Image-based profiling approaches, including spectral-unmixing and cell painting, have shown that organelle morphology and organization encode information about cellular identity, stress responses, and disease-associated phenotypes (*32–34*). However, these approaches were developed for cultured cells and are difficult to extend to intact tissues, where complex three-dimensional geometry and cellular diversity present major challenges for organelle segmentation, feature extraction, and spatial interpretation. As a result, scalable strategies for quantifying multi-organelle architecture across cells within native tissue environments remain limited.

Here we present *spatial Organellomics* (*sOrganellomics*), an imaging and analysis workflow that quantifies multi-organelle architecture at single-cell resolution in intact tissues. By combining multiplexed three-dimensional fluorescence imaging, automated segmentation, and interpretable feature extraction, *sOrganellomics* generates high-dimensional structural signatures from millions of organelles across thousands of individual cells. This approach provides an interpretable structural fingerprint of cell state that complements molecular profiling approaches and enables systematic analysis of functional cellular diversity *in situ*.

We first applied *sOrganellomics* across metabolically specialized organs and found that organelle architecture distinguishes major cellular identities in the pancreas and liver. We then used hepatocytes in the liver as a model system to test whether organelle architecture could resolve functional diversity within a single cell type, taking advantage of the well-established metabolic zonation along the hepatic lobule (*35–37*). Using this system, we found that hepatocyte diversity is not organized as graded homogeneous layers along the zonation axis, as proposed by classical models of liver zonation, but instead distinct hepatocytes form intermixed communities that generate a previously unrecognized sub-zonal diversity organization (*35–37*). We further show that fasting and Western diet differentially remodel this organization. Importantly, intravital imaging experiments demonstrate that these organelle architectural changes are associated with altered mitochondrial activity *in vivo*. Multi-organelle signatures also classify nutritional conditions and predict the progression of early metabolic disease. Together, these findings establish organelle architecture as a scalable and interpretable dimension of cell state and tissue adaptation.

## Results

### sOrganellomics distinguishes cellular identities across organs through organelle architecture

Cellular states emerge from regulatory programs that shape the internal organization of the cell (Fig. 1A). Organelle architecture—defined here by the morphology, abundance, and spatial distribution of intracellular compartments—reflects the metabolic and functional demands imposed by the cellular microenvironment (*16*, *18*, *19*, *27–31*, *38*, *39*). To quantify this organization systematically in intact tissues, we developed *spatial Organellomics* (*sOrganellomics*), a microscopy-based workflow that combines multiplexed three-dimensional fluorescence imaging via confocal microscopy, automated Cellpose-based segmentation, and feature extraction to generate single-cell structural profiles from tissue sections (Fig. 1A–B, fig. S1) (*40*). In this study, we focused on mitochondria, peroxisomes, and lipid droplets because these organelles are closely linked to redox balance, bioenergetics, and lipid metabolism in many cell types (*28–31*, *41–44*). However, *sOrganellomics* is not tied to a fixed marker panel; organelle selection can be adapted to the tissue and biological question. For each segmented cell, *sOrganellomics* extracts approximately 100 interpretable features describing organelle abundance, morphology, and intracellular distribution, yielding high-dimensional organelle signatures capturing the architecture of millions of organelles across thousands of cells *in situ* (fig. S1).

**Fig. 1.**
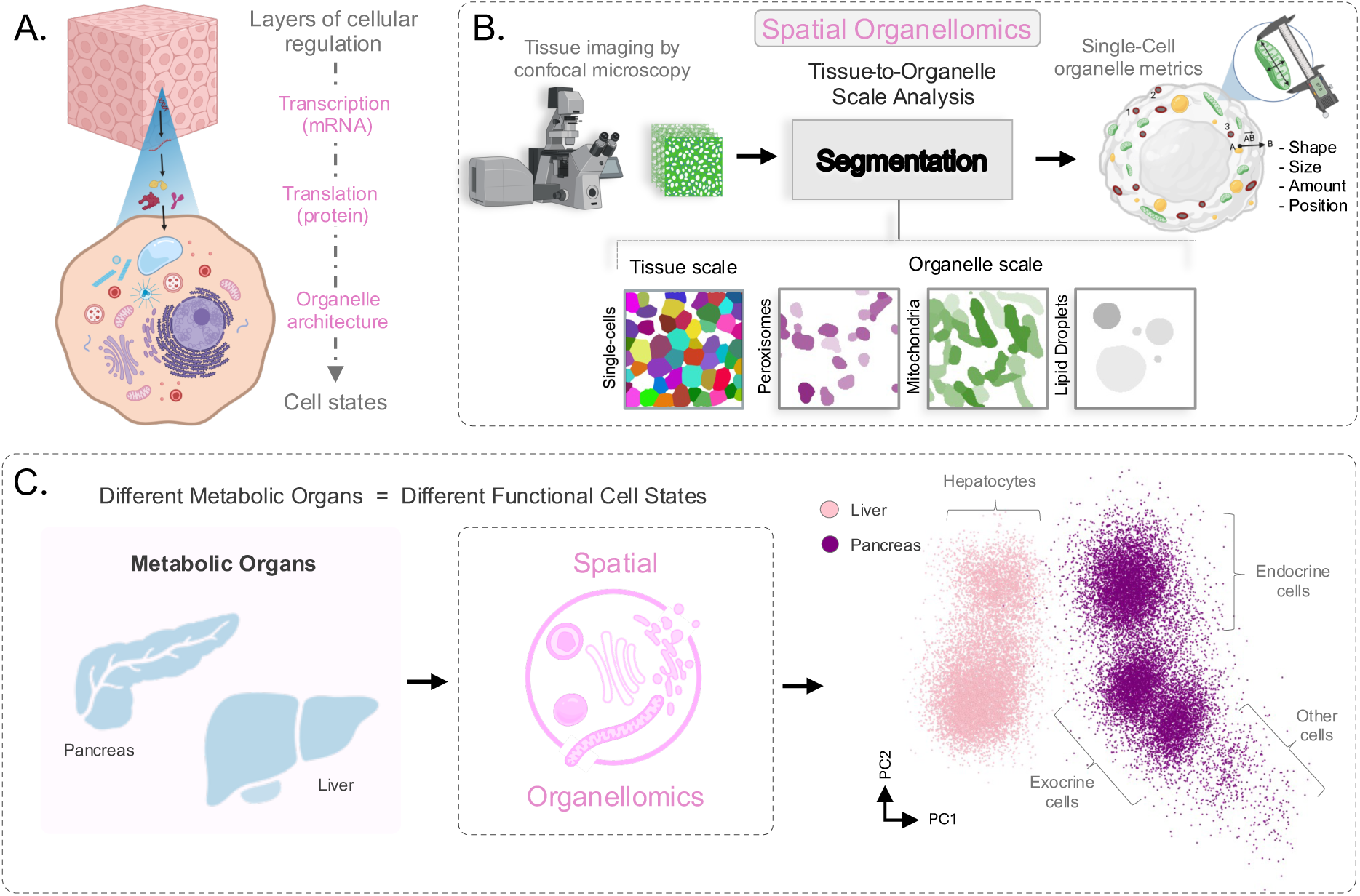
*sOrganellomics* quantifies multi-organelle architecture to define cell states across metabolic organs. **(A)** Schematic illustrating how multiple layers of cellular regulation shape organelle architecture, reflecting underlying cellular states. **(B)** Overview of the *sOrganellomics* workflow. Confocal images of tissue sections labeled for cell boundaries, mitochondria, peroxisomes, and lipid droplets were analyzed using custom instance-segmentation models to identify cells and organelles, followed by extraction of quantitative single-organelle features from the segmented masks. Representative segmentation outputs at tissue (cell) and intracellular (organelle) scales are shown. **(C)** Schematic overview of the workflow from liver and pancreas tissue through *sOrganellomics* to downstream analysis by principal component analysis (PCA). PCA of single cells derived from liver (*n* = 4 mice; >4000 cells) and pancreas (*n* = 3 mice; >8000 cells). Each point represents one cell. Liver and pancreas cells separate in PCA space based on organelle features, and within pancreas samples, endocrine, exocrine and other cell populations form distinguishable subpopulations. Endocrine cells were identified by insulin and glucagon labeling, whereas exocrine cells were defined by their characteristic size, morphology and absence of insulin and glucagon signal; cells not meeting these criteria were classified as other.

To test the premise that organelle architecture encodes information to define cell state, we first asked whether multi-organelle signatures could distinguish cellular identities across metabolically specialized organs. We applied *sOrganellomics* to the liver and the pancreas generating organelle signatures for more than 12,000 cells. Dimensionality reduction of these features using principal component analysis (PCA) revealed clear segregation of cells according to both tissue of origin and cellular identity (Fig. 1C). Liver hepatocytes formed a distinct point-cloud separated from pancreatic cells, while pancreatic endocrine, exocrine, and other cell populations further segregated based on organelle architecture alone. These results indicate that multi-organelle signatures provide an alternative layer of information to resolve cellular populations across organs, establishing *sOrganellomics* as a scalable approach for defining cellular identity from subcellular organization *in situ*.

### Organelle architecture resolves hepatocyte heterogeneity beyond molecular profiling

Having established that organelle architecture distinguishes cellular populations across organs, we next asked whether it could resolve functional diversity within a single cell type. This is a more demanding test because cells of the same lineage can perform distinct metabolic functions despite sharing broad transcriptional identity (*27*, *45–47*). The liver provides an ideal model system for this question because hepatocytes are exposed to gradients of oxygen, nutrients, and hormones along the portal vein (PV)-to-central vein (CV) axis, giving rise to spatially organized metabolic specialization across the hepatic lobule (Fig. 2A) (*35–37*, *48*, *49*).

**Fig. 2.**
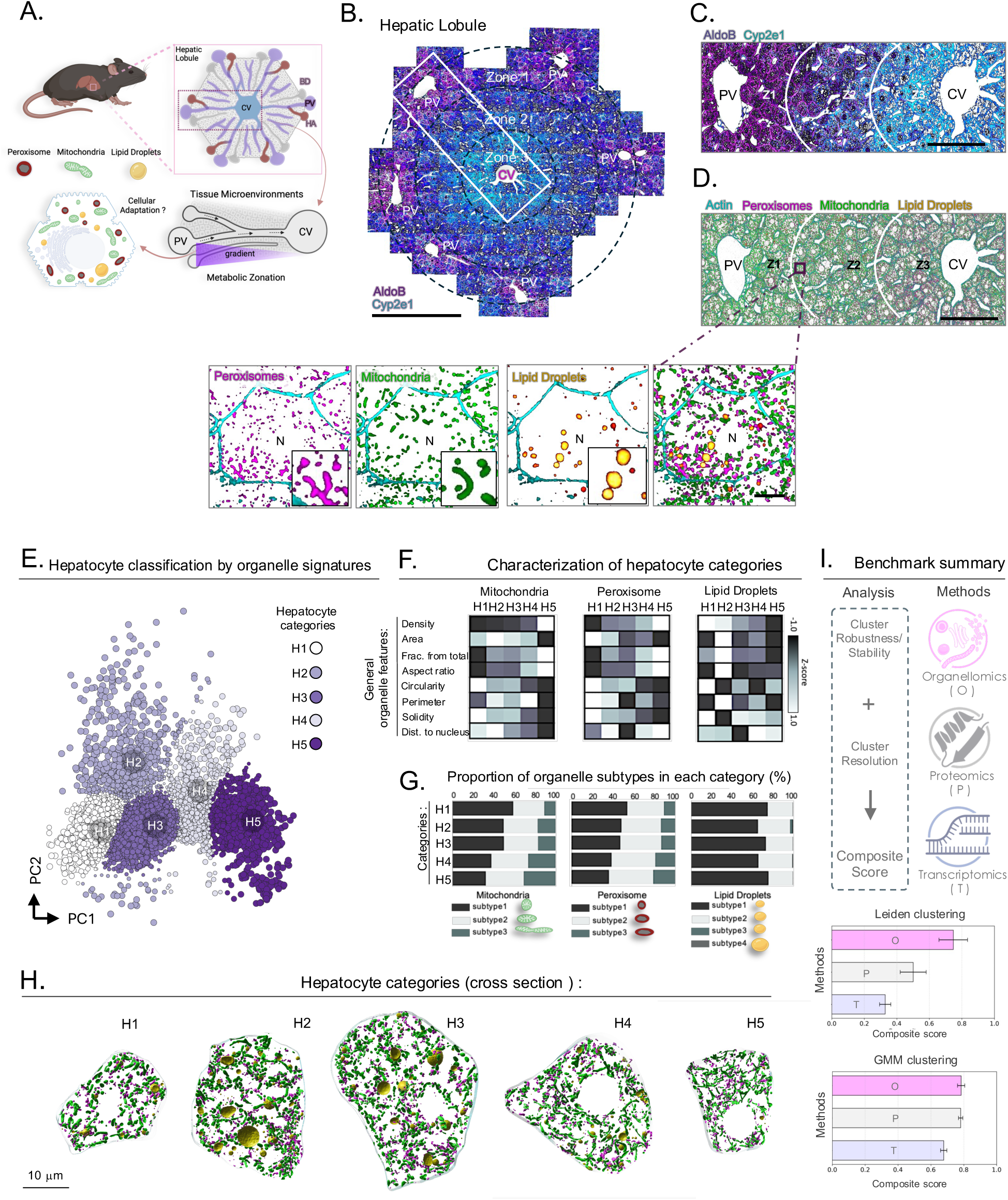
Single-cell mapping of organelle architecture reveals hepatocyte heterogeneity beyond molecular profiling. **(A)** Schematic of the liver lobule and linearized acinus illustrating gradients of nutrients, oxygen, and hormones from the portal vein (PV) to the central vein (CV). HA, hepatic artery; BD, Bile duct. **(B)** Confocal image of a representative liver lobule showing PVs at the periphery and a CV at the center. Zonation markers Aldolase B (AldoB; periportal enriched) and CYP2E1 (pericentral enriched) delineate zones 1–3. An acinus is marked by a white dashed box. Scale bar, 300 μm. **(C)** Inset from B showing a magnified crop of the acinus spanning from PV to CV including all three liver zones (Z1, Z2, Z3). Scale bar, 100 μm. **(D)** Representative fluorescence image of an acinus from control-fed mice showing hepatocytes, peroxisomes, mitochondria, and lipid droplets (LDs) across the PV–CV axis. Insets highlight single-cell and single organelle resolution. Scale bars, 100 μm and 5 μm (inset). N = position of the nucleus. **(E)** PCA visualization of single hepatocytes from control livers (4 mice; ∼3 lobules per mouse, ∼3 acini per lobule; >4000 hepatocytes). Clustering was performed with PCs that detected most stable clusters using a multi-metric Gaussian mixture model (GMM) approach. Five hepatocyte categories (H1–H5) were identified (see Methods). **(F)** Heatmaps of normalized organelle features across hepatocyte categories showing distinct multi-organelle architectural signatures. **(G)** Distribution of organelle subtypes within hepatocyte categories. Mitochondria and peroxisomes were classified by aspect ratio and LDs by area. **(H)** Representative 3D reconstruction of hepatocyte categories H1–H5. **(I)** Benchmarking of spatial Organellomics (O), spatial proteomics (P), and spatial transcriptomics (T) using matched features (98) and cell budgets (390). Clustering quality was evaluated with Leiden clustering and GMMs using multiple metrics scaled to [0,1] and combined into a composite clustering score (maximum = 1). Bars represent mean ± bootstrap variability of computed composite score.

To investigate hepatocyte functional diversity *in situ*, we applied *sOrganellomics* to mouse liver sections spanning the hepatic lobule (Fig. 2B). In addition to mitochondria, peroxisomes, lipid droplets, and cell boundaries, we stained for Aldolase B and CYP2E1 to mark periportal and pericentral regions and confirm that the full PV–CV axis was captured (Fig. 2C-D, Movie S1 and Movie S2) (*50–52*). For each segmented hepatocyte, we quantified organelle features together with spatial position, enabling systematic analysis of intracellular organization across the lobule (figs. S2–S7).

Dimensionality reduction of organelle features (PC1: 32.2% variance, PC2: 11.1% variance) followed by an unsupervised multi-metric clustering strategy via Gaussian Mixture Models (GMM) revealed five stable hepatocyte categories (H1-H5) that were reproducible across control livers (Fig. 2E, fig. S8) (*53–56*). Although additional PCs captured more variance, cluster stability declined beyond PC2 (fig. S8C). These categories differed in mitochondrial morphology, peroxisome abundance, and lipid droplet size distributions, reflecting distinct intracellular organizations driven by variations in organelle subtypes (Fig. 2F-H, fig. S9-S11, Movie S3-S8). For example, H1 hepatocytes contained a greater proportion of larger, rounder mitochondria positioned closer to the cell center, whereas H5 hepatocytes showed the opposite trend (Fig. 2F-H, Movie S4 and Movie S8). Three-dimensional reconstructions highlighted the extent of these architectural differences across categories (Fig. 2H). One-way ANOVA confirmed significant differences in organelle features between hepatocyte categories, with the most notable differences observed between H5 and the other categories (fig. S8G). Together, these data indicate that multi-organelle architecture resolves reproducible hepatocyte states within the liver.

To assess how well organelle-based profiling captures hepatocyte heterogeneity relative to other spatial modalities, we compared *sOrganellomics* with published single-cell spatial transcriptomic and spatial proteomic liver datasets using a harmonized analysis (*57*, *58*). After matching cell numbers and feature dimensions across datasets, clustering performance was evaluated using both Leiden clustering and Gaussian mixture modeling to reduce method-specific bias. Across metrics capturing cluster robustness, separation, and resolution, *sOrganellomics* performed strongly and was comparable to, or better than, the spatial molecular datasets included in this benchmark (Fig. 2I, fig. S12A, see Methods). These comparisons support the idea that organelle architecture provides a complementary and high-resolution dimension for resolving hepatocyte heterogeneity *in situ*.

### sOrganellomics reveals that hepatocyte states are not fully explained by zonal position

Historically, models of liver zonation have proposed that hepatocytes adopt distinct metabolic roles primarily according to their position along the PV–CV axis, resulting in layered zones with relatively homogeneous cellular functions within each zone (*35–37*, *48*, *49*). Having identified five hepatocyte states from organelle architecture, we next asked whether these states simply reflected position along this axis or instead captured additional structure within the hepatic organization (Fig. 3A).

**Fig. 3.**
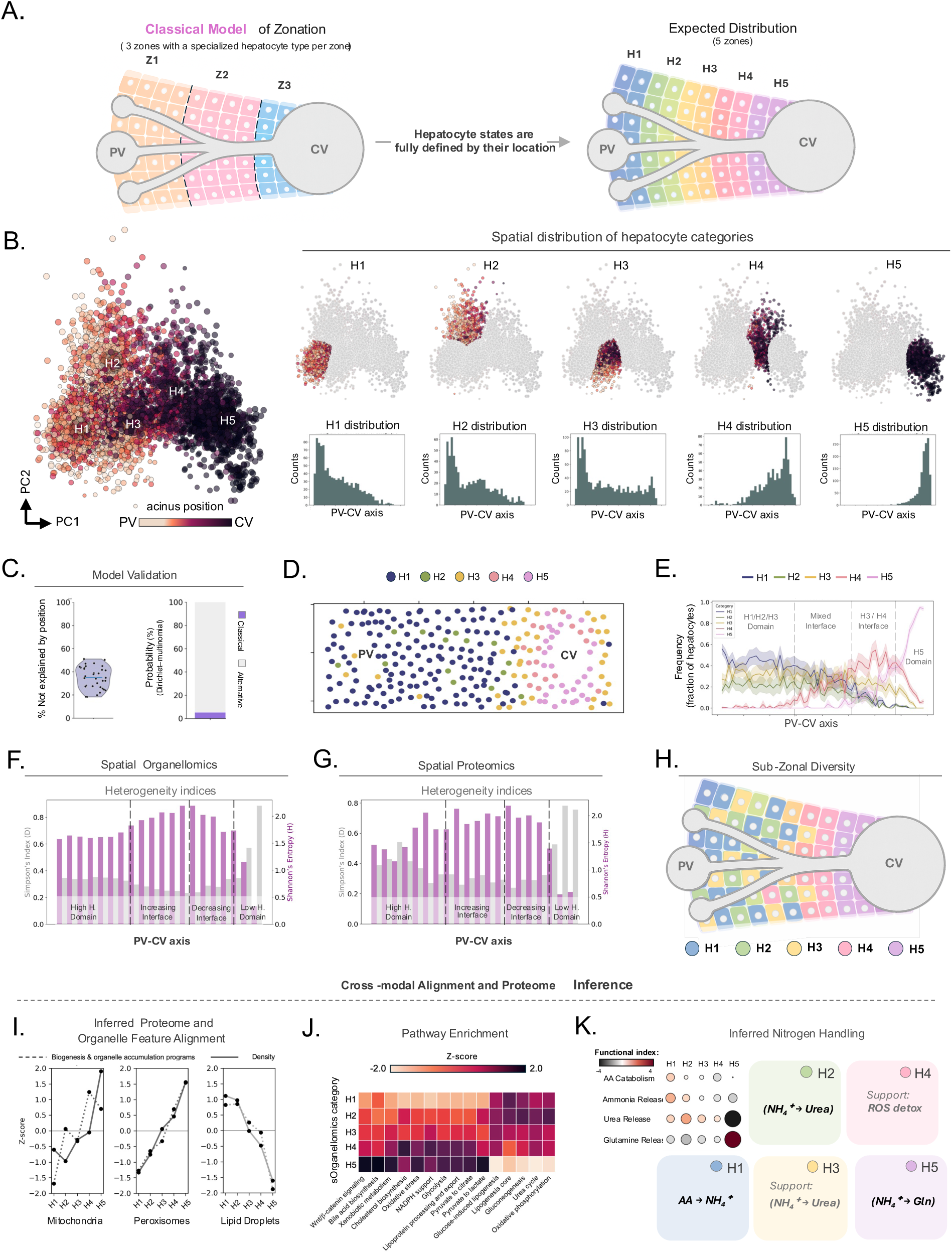
*sOrganellomics* reveals sub-zonal diversity and putative metabolic specialization of hepatocyte states. **(A)** Schematic of the classical model of liver zonation with three zones along the PV–CV axis, each containing homogeneous hepatocyte populations that are distinct between zones and in which cell identities are strictly determined by spatial position (left). Based on this model, the expected distribution of five hepatocyte categories (H1–H5) is shown (right). **(B)** PCA visualization of hepatocytes from control livers (n = 4 mice; >4000 cells). Colors indicate normalized PV–CV position (white, periportal; purple, pericentral). The histograms show distributions of hepatocyte categories (H1–H5) across the PV–CV axis. **(C)** Validation of hepatocyte category distribution in an acinus based on the classical model of liver zonation. The violin plot shows percentage of hepatocyte-category variability not explained by position across acini (n=30 acini). The stack bar shows posterior probabilities of position-dependent versus alternative generative models computed per acinus using a Dirichlet–multinomial framework (see Methods). **(D)** Representative acinus showing spatial coexistence of hepatocyte states. **(E)** Spatial frequency profiles of hepatocyte categories along the PV–CV axis (mean ± SEM; n = 30 acini from 4 mice). **(F, G)** Pittsburgh Heterogeneity Index analysis showing Simpson’s dominance (D) and Shannon entropy (H) of hepatocyte-category distributions along the PV–CV axis derived from *sOrganellomics* (F) and spatial proteomics (G). **(H)** Revised model of sub-zonal liver organization based on the intermixed spatial distribution of organelle-defined hepatocyte states. **(I–K)** Cross-modal alignment linking organelle-defined hepatocyte states to proteomic programs from independent spatial proteomics (see Methods). (I) Comparison of inferred organelle-related programs with corresponding imaging-derived organelle measurements. Inferred mitochondrial, peroxisomal, and lipid droplet biogenesis and accumulation programs were compared with corresponding imaging-based organelle densities across hepatocyte states. (J) Heatmap showing z-scored enrichment of selected metabolic pathways across hepatocyte states. (K) Example of inferred metabolic pathway activity based on nitrogen handling balance across hepatocyte states. Dot size and color indicate the relative magnitude and direction of the functional bias (z-scored across categories). Distinct nitrogen-handling profiles include amino acid (AA)–derived ammonia production (H1), balanced urea routing (H2–H3), glutamine (Gln) synthesis bias (H5), and oxidative stress (ROS)–associated metabolism (H4).

Surprisingly, mapping hepatocyte categories across the hepatic acinus showed that most states were distributed across broad portions of the lobule, coexisting in different proportions rather than confined to graded homogeneous layers (Fig. 3A–E, fig. S12B). Some categories were enriched near periportal regions (H1, H2, and H3), whereas a single category was preferentially associated with pericentral areas (H5). In addition, other categories (H3 and H4) frequently occurred at the boundaries between neighboring domains, forming transition interfaces across the acinus (Fig. 3D–E). Because clustering was performed using organelle features alone, we next asked to what extent PV–CV position accounted for the observed category structure. Position-only models did not fully explain this organization: residual spatial autocorrelation, position-based classification performance, and the fraction of category variability explained by position all indicated that lobular location captured only part of hepatocyte diversity, leaving about 40% of category variability unexplained (Fig. 3C, fig. S13) (*59–61*). These findings indicate that hepatocyte categories are influenced by zonal position but are not fully determined by it.

Spatial Pittsburgh Heterogeneity Index analysis further revealed extensive mixing of hepatocyte categories across the acinus. The highest diversity occurred within interface regions of the lobule, whereas specific states became more dominant toward the central vein (H5) (Fig. 3F) (*62*, *63*). To compare organelle-derived spatial heterogeneity with molecular measurements, we applied the same spatial heterogeneity analyses to a published single-cell spatial proteomics dataset from the mouse liver (*58*). The proteomic data recapitulated similar PV–CV heterogeneity trends, broadly matching those observed by *sOrganellomics* (Fig. 3F–G, fig. S12B-D). Organelle-based analysis, however, resolved sharper local interfaces between domains and finer spatial transitions, consistent with its denser cellular sampling. This suggests that intracellular architecture captures spatial organization consistent with proteomic heterogeneity while providing higher resolution for defining the cellular organization within the tissue. Furthermore, these findings support a refined model of liver organization in which hepatocytes form intermixed communities within each zone generating a sub-zonal diversity organization across the hepatic acinus (Fig. 3H). To test whether organelle-defined states were associated with underlying molecular programs, we then aligned the imaging and proteomic datasets using a probabilistic cross-modal framework at the hepatocyte-category level (fig. S14, see Methods). Enrichment analyses after alignment showed that organelle-related molecular programs mirrored the structural features measured by *sOrganellomics*. Inferred proteomic programs associated with mitochondrial, peroxisomal, and lipid droplet abundance closely matched the corresponding organelle measurements from imaging, consistent with a relationship between organelle-defined categories and underlying proteomic programs (Fig. 3I). Inferred pathway enrichments further suggested that hepatocyte categories differ in glycolysis, gluconeogenesis, amino acid metabolism, oxidative stress responses, and lipid handling (Fig. 3J). In this framework, neighboring hepatocyte categories may participate in related, and in some cases potentially complementary, metabolic programs within shared spatial domains, including pathways involved in nitrogen handling (Fig. 3K).

Together, these findings demonstrate that hepatocyte diversity is not organized as graded layers of homogeneous hepatocyte states defined solely by position as previously thought (*35–37*, *48*, *49*). Instead, *sOrganellomics* reveals a sub-zonal diversity organization in which intermixed hepatocyte states within each zone may collectively coordinate metabolic tasks. Integration with spatial proteomics further suggests that neighboring hepatocyte states perform complementary steps within shared metabolic pathways, supporting a model for how hepatocyte communities distribute metabolic labor within the liver. We next asked whether this organization is remodeled in response to nutritional stress.

### Nutritional perturbations differentially remodel the sub-zonal hepatocyte diversity

Nutritional conditions strongly influence tissue metabolism by altering cellular bioenergetic demands (*64*, *65*). Having established that hepatocytes form a sub-zonal diversity organization, we next asked whether this organization represents a fixed anatomical pattern or an adaptive feature of liver physiology. We reasoned that, if it is adaptive, then acute and chronic metabolic perturbations should remodel it in distinct ways. To address this question, we applied *sOrganellomics* to liver sections from mice subjected to three nutritional conditions: control feeding, overnight fasting, and Western diet (WD) feeding for one month (Fig. 4A-C).

**Fig. 4.**
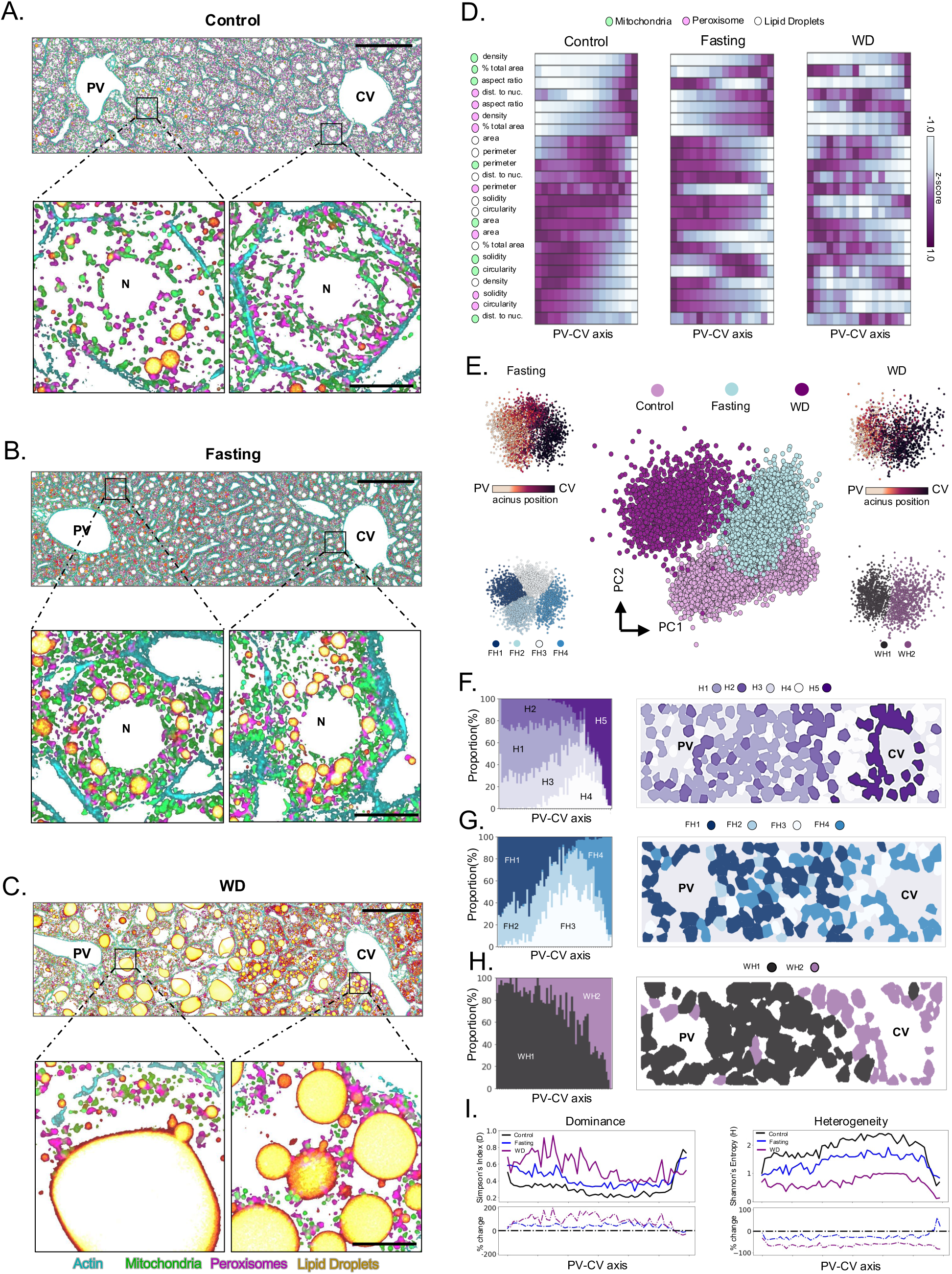
Nutritional perturbations remodel sub-zonal hepatocyte heterogeneity across the PV– CV axis. (A–C) Representative confocal images of liver acini from control-fed **(A),** overnight-fasted **(B)**, and 4-weeks western diet (WD)-fed **(C)** mice following fluorescent labeling of cell boundaries and organelles as labeled. Insets show higher magnification views of periportal (PV) and pericentral (CV) regions. Scale bars, 100 μm; inset, 5 μm. **(D)** Spatial heatmaps of selected organelle features across the normalized PV–CV axis for control (n=4 mice), fasting (n=3 mice), and WD (n=3 mice) groups. Colors represent z-scored feature values within each condition. **(E)** PCA visualization of hepatocytes (∼10,500 cells total) from control, fasting, and WD groups; separation between nutritional groups is evident in the PCA space. Sub-clustering within fasting and WD groups was performed using multi-metric GMM, identifying four fasting categories (FH1 to FH4) and two WD categories (WH1 and WH2). Categories are color-coded as indicated. Spatial distributions of these subclusters along the PV–CV axis are shown above each subpanel. **(F–H)** Proportion (left) and PV–CV spatial distribution (right) of hepatocyte categories along PV-CV axis in control **(F)**, fasting **(G)**, and WD **(H)** groups. **(I)** Pittsburgh Heterogeneity Index analysis showing Simpson’s dominance (D) and Shannon entropy (H) across the PV–CV axis for each nutritional condition. Lower panels show the percent change in heterogeneity in fasting and WD groups compared to controls.

Both fasting and WD feeding induced pronounced remodeling of organelle architecture relative to controls (Fig. 4A–D, Movie S9). Consistent with increased metabolic demand during fasting (*66*, *67*), hepatocytes exhibited higher mitochondrial and peroxisomal densities with a greater proportion of elongated subtypes, along with a global increase in lipid droplets likely reflecting lipid mobilization from adipose tissue (*68*, *69*) (fig. S15–S18). In contrast, WD feeding reduced peroxisome abundance, maintained mitochondrial density but shifted towards rounded subtypes, and increased lipid droplet accumulation by approximately fivefold, with large droplets particularly enriched near periportal regions (fig. S18). Together, these findings highlight distinct organelle adaptation strategies in hepatocytes in response to nutrient deprivation versus nutrient excess (*70*, *71*).

We next examined how these changes influence hepatocyte states across nutritional conditions. PCA revealed that hepatocytes segregated according to nutritional state (Fig. 4E). Within each condition, GMM clustering revealed a reduction in hepatocyte categories in fasted (4 categories) and WD fed (2 categories) animals compared to controls (5 categories), yielding a total of 11 distinct categories across all groups (fig. S19). Thus, metabolic perturbation did not simply shift existing hepatocyte categories along the acinus, but instead reshaped the landscape of organelle configurations to generate new hepatocyte states.

Despite differences in the number of hepatocyte categories under each condition, multiple states continued to coexist to some extent across the PV–CV axis (Fig. 4F–H, fig. S19A–F). However, both fasting and WD feeding reduced overall heterogeneity along the lobule, with the most pronounced reduction observed after WD feeding. This condition was also associated with greater state dominance in Pittsburgh Heterogeneity Index analyses, whereas fasting selectively increased heterogeneity near the central vein region (Fig. 4I).

*sOrganellomics* thus reveals that sub-zonal hepatocyte diversity is not a fixed feature of liver organization, but a dynamic one that is remodeled by nutritional perturbation. These adaptive responses reshape organelle architecture, generating new hepatocyte states while altering the overall degree of diversity across the lobule. Notably, acute metabolic stress induced by fasting largely preserved sub-zonal diversity, whereas chronic nutrient excess under WD conditions led to a pronounced loss of this organization.

### Intravital imaging links fasting-associated architectural remodeling to altered mitochondrial activity in vivo

The relationship between control and fasting hepatocyte states suggested that acute metabolic adaptation might follow an ordered remodeling trajectory. In PCA space, WD hepatocytes occupied a distinct architectural region with limited overlap with controls, consistent with more extensive reorganization under chronic nutrient excess (Fig. 4E). By contrast, hepatocytes from overnight-fasted mice remained topologically connected to control states despite forming distinct fasting-associated categories, suggesting a more continuous structural transition during acute adaptation.

To examine this possibility, we applied probabilistic trajectory inference to hepatocyte organelle signatures. The inferred transition landscape was non-random and suggested convergence of control hepatocytes onto an intermediate state before divergence into fasting-associated states. The highest-probability path proceeded through a transition hub (H4-to-FH2) and subsequently distributed across fasting configurations, indicating an ordered organelle remodeling (Fig. 5A-B, fig. S20). The dominant remodeling axis (PC1) was defined primarily by coordinated changes in mitochondrial subtype composition and abundance, increased peroxisome content, and lipid droplet expansion (fig. S20). Progression along this axis converged with global transition topology (ρ ≈ 0.83), indicating that the principal axis of organelle remodeling closely tracked the inferred shift from control to fasting-associated states (Fig. 5B). When projected back into tissue space, this axis showed stronger remodeling in hepatocytes near the CV than near the PV (Fig. 5C), suggesting region-specific adaptation to fasting.

**Fig. 5.**
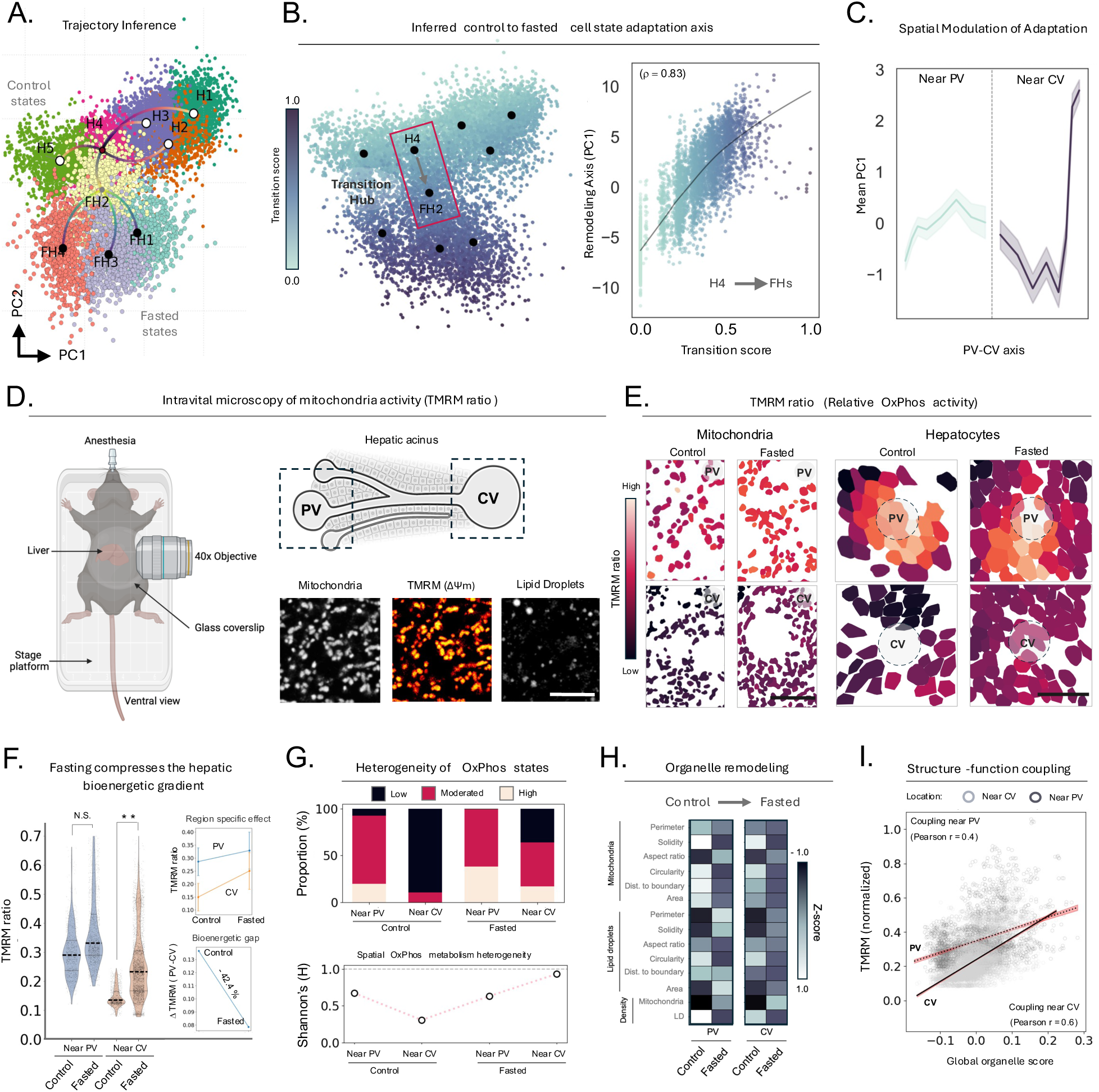
Organelle remodeling during fasting links structural state transitions to functional adaptation *in vivo*. **(A)** PCA visualization of hepatocyte categories from control and fasted livers, highlighting the most probable transition trajectories from control to fasting states. Points represent individual cells colored by category (control: H1–H5; fasting: FH1–FH4). White dots indicate centroids of control categories, and black dots indicate centroids of fasting categories. The centroid of control H4 is shown in red, and the centroid of FH2 is shown in gold. **(B**) Visualization of probabilistic trajectory analysis in PCA space, showing an intermediate transition hub between H4 and FH2 states (red dashed box). A corresponding plot shows a strong correlation between projection onto the dominant remodeling axis (PC1) and control-to-fasted transition score (Spearman ρ ≈ 0.83). **(C)** Mean PC1 plotted along the normalized PV–CV axis (mean ± SEM) shows spatial bias in organelle adaptation during control to fasted states transition. **(D)** Intravital microscopy (IVM) setup for *in vivo* imaging and acinus schematic indicating imaging regions near PV and CV (dashed boxes). Representative images show mitochondria, TMRM (mitochondrial membrane potential, ΔΨm), and lipid droplets. **(E)** TMRM ratio (TMRM/mito-Dendra2) at subcellular and cellular level in periportal (PV) and pericentral (CV) regions of control and fasted groups. Statistics were derived from a linear mixed-effects model fit to 9,585 cells from 6 livers, with liver and tile included as random effects. **(F)** Quantification of mitochondrial membrane potential under control and fasted conditions, shown as violin plots with quartiles indicated by dotted lines and raw cell-level distributions in black points. PV did not differ significantly between control and fasted samples (p = 0.346; N.S.), whereas CV increased after fasting (p = 0.0259; **). The top inset shows the mixed-model mean TMRM ratio values ± 95% CI in PV and CV across experimental groups, while the bottom inset shows the mixed-model estimated difference in TMRM between hepatocytes near PV and those in CV (TMRM ratio in PV − TMRM ratio in CV, or Bioenergetic gap), indicating a 42.4% gap reduction in fasting conditions (p = 1.17 × 10⁻²⁴; ***). **(G)** Proportions of TMRM-defined bioenergetic states derived from Gaussian mixture modeling of normalized TMRM intensity (top) and corresponding Shannon entropy quantifying OxPhos state heterogeneity (bottom). **(H)** Heatmap of control-to-fasting organelle feature remodeling in PV and CV regions. The color of the scale on the right indicates z-scored feature values. **(I)** Structure to function coupling of TMRM intensity and a global organelle score (see Methods) derived from intravital organelle features. Lines indicate ordinary least squares fit with 95% confidence intervals. Stronger slope and higher correlation in CV-hepatocytes (Pearson r = 0.6, R^2^ = 0.36) as compared to PV-hepatocytes (Pearson r = 0.4, R^2^ = 0.15) indicate enhanced architecture–activity coupling near the CV. Significance: N.S., p > 0.05; **, 0.05 ≥ p > 0.0001; ***, p ≤ 0.0001.

We then asked whether this spatial bias corresponded to functional metabolic changes *in vivo*. Using intravital microscopy, we measured mitochondrial morphology, LD features, and mitochondrial membrane potential in periportal and pericentral hepatocytes using TMRM-based imaging (Fig. 5D) (*72–74*). Fasting increased TMRM intensity ratios across the lobule, consistent with increased mitochondrial activity interpreted here in the context of enhanced oxidative phosphorylation (OxPhos) (Fig. 5E-F) (*75*). However, these changes were disproportionately greater in hepatocytes near the CV. As a result, the basal PV–CV bioenergetic gradient observed in control livers was reduced by 42.4% after fasting (Fig. 5F). Gaussian mixture modeling of single-cell TMRM ratios identified three hepatocyte groups with low, intermediate, and high mitochondrial activity states. Under control conditions, hepatocytes near the PV displayed broader metabolic heterogeneity, whereas hepatocytes near the CV were more homogeneous strongly enriched for low-activity states consistent with sub-zonal functional diversity *in vivo* (Fig. 5G). Fasting modestly shifted hepatocytes near the PV toward higher-activity states, but more substantially redistributed hepatocytes near the CV from the low-activity group into intermediate and high-activity states, increasing functional heterogeneity in that region (Fig. 5G). These metabolic changes paralleled the corresponding architectural remodeling: fasting-associated shifts in mitochondrial morphology and LD expansion were coupled to TMRM measurements, with stronger structure-function coupling in hepatocytes near the CV (Fig. 5H–I). Together, these results indicate that acute fasting induces a spatially biased trajectory of organelle remodeling that enhances cellular activity and demonstrate that these structural adaptations correspond to functional metabolic shifts, linking organelle architecture to metabolic state transitions *in vivo*.

### Multi-organelle signatures predict nutritional status and early disease progression

Having established that organelle architecture tracks hepatocyte state and metabolic adaptation, we next asked whether these structural signatures could be used predictively. To test this, we used Multilayer Perceptron (MLP) models using organelle metrics as descriptors (*76*, *77*). We randomized our datasets and used 80% of all experimental groups for training and 20% for validation of our models (Fig. 6A).

**Fig. 6.**
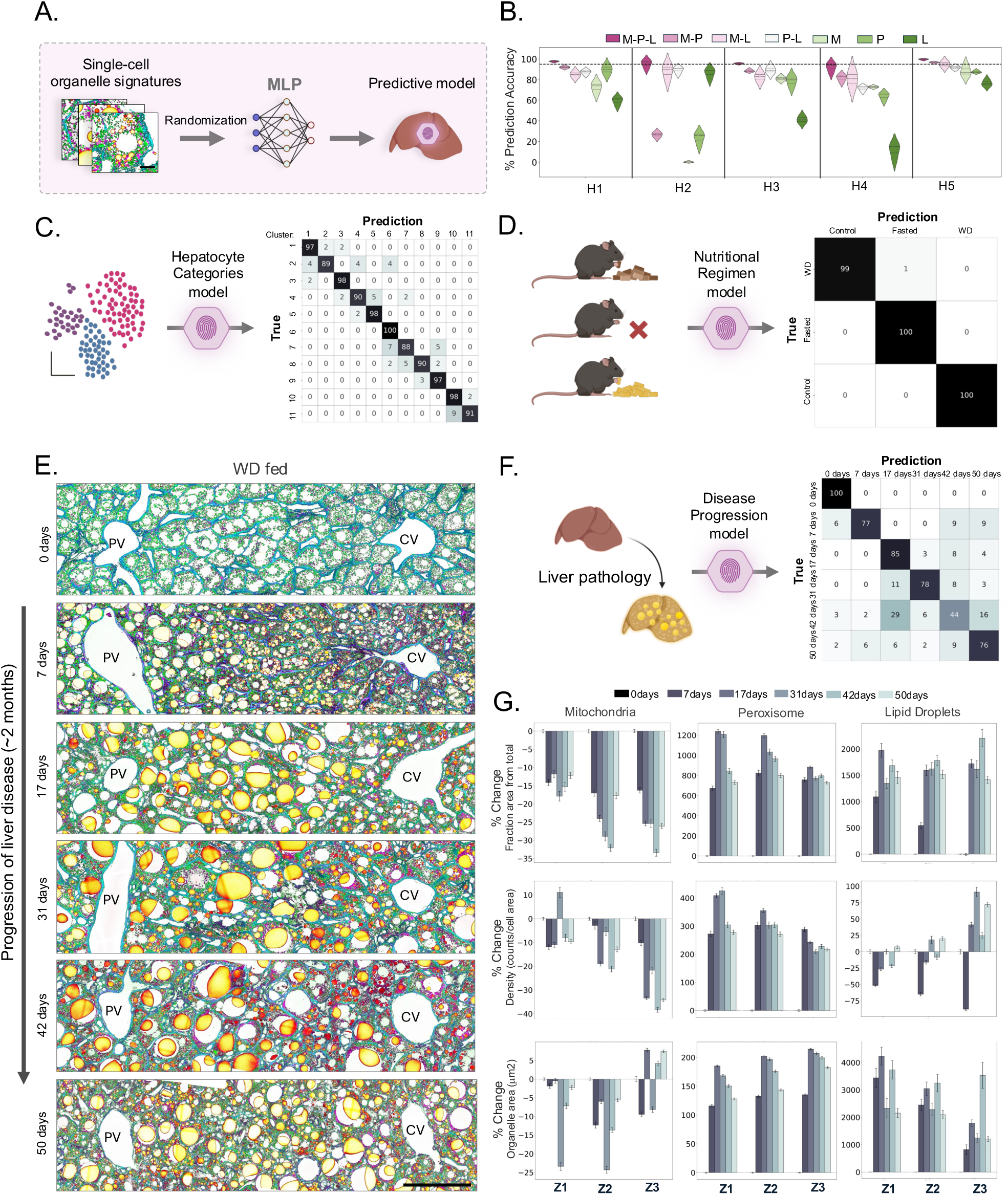
Organelle signatures enable predictive modeling of hepatocyte states and disease progression. **(A)** Schematic of the multilayer perceptron (MLP) architecture used to predict cellular states from single-cell organelle signatures. The model consisted of three hidden layers (64 neurons each). Training and validation were performed with 80% and 20% of total data respectively (see Methods). Separate models were trained to predict hepatocyte category, nutritional regimen, and disease progression stage. **(B)** Violin plots illustrate the contribution of mitochondria (M), peroxisomes (P), and lipid droplets (L), individually or in combination (as indicated), to predict hepatocyte categories from the control group. **(C)** Confusion matrix for validation of hepatocyte category prediction across all nutritional groups (11 total categories). Values represent percentage of correct predictions within the validation set. **(D)** Confusion matrix for prediction of nutritional regimen. Values represent percentage of correct predictions within the validation set. **(E)** Representative liver acini from western diet (WD)-fed mice at the indicated time points (0–50 days). Scale bar, 100 μm. **(F)** Confusion matrix for prediction of liver disease stage in WD-fed mice. Values represent percentage of correct predictions within the validation set. **(G)** Percentage change in organelle features (mitochondria, peroxisomes, lipid droplets) within each classical liver zone of WD-fed mice relative to control (0 days). Bars represent mean ± SEM across acini within each time point (total = 21,422 cells).

We first tested whether organelle signatures could predict distinct hepatocyte states in control livers. Models trained on features from individual organelles, organelle pairs, or all three organelles showed that multi-organelle signatures produced the highest overall performance, yielding approximately 95% mean classification accuracy across hepatocyte categories (Fig. 6B). Some categories, particularly H5 hepatocytes, were classified accurately even from single-organelle or paired-organelle features (80–96%), consistent with their distinct structural profiles. Similar trends were obtained with logistic regression and random forest classifiers, indicating that the signal was not specific to one model class (fig. S21).

Extending this approach across all experimental groups, including control, fasting, and WD, maintained similarly high performance (∼95% average accuracy) (Fig. 6C). Strikingly, nutritional condition was predicted with near-perfect accuracy (∼99%) (Fig. 6D), supporting the idea that organelle architecture constitutes a robust structural fingerprint defining cell state diversity and nutritional status.

Encouraged by these results, we investigated whether *sOrganellomics* could help predict subcellular changes associated with metabolic disease progression. As a model of early metabolic dysfunction-associated steatotic liver disease (MASLD), male mice were fed WD for up to 50 days and liver samples collected at multiple time points (*78*). Signs of steatosis, including progressive lipid accumulation accompanied by changes in mitochondrial and peroxisomal morphology, became apparent by day 7 and intensified over time, ultimately leading to substantial reorganization of tissue architecture (Fig. 6E) (*79*, *80*). An MLP model trained to classify disease stage from organelle features achieved approximately 80% mean accuracy across time points (Fig. 6F). These stage-dependent differences were associated with dynamic redistributions of organelle features, including temporal and zone-specific remodeling across disease progression (Fig. 6G). This suggests that progressive pathological remodeling is encoded within organelle architectural changes.

These findings not only underscore the sensitivity of organelle signatures as indicators of metabolic transitions, but also provide a predictive “structural code” for cellular state, nutritional status, and disease progression. More broadly, this approach provides a workflow for tracking organelle-level pathological changes in tissues using off-the-shelf machine learning methods and may have significant implications for understanding later-stage diseases in the liver or other metabolic organs.

## Discussion

Understanding how tissues organize and adapt diverse cellular states remains a central challenge in cell and tissue biology. Although transcriptomic approaches have transformed efforts to define cell identity, they do not directly capture the intracellular organization through which many cellular functions are executed and often sacrifice spatial and structural context (*52*, *81*, *82*). Spatial transcriptomic and proteomic methods have begun to bridge this gap, but they often remain limited in throughput, subcellular resolution, or accessibility (*58*, *83*, *84*). Here we introduce *spatial Organellomics* (*sOrganellomics*), an imaging-based workflow for quantifying multi-organelle architecture at single-cell resolution in intact tissues. By combining multiplexed three-dimensional fluorescence imaging with automated segmentation and interpretable feature extraction, this approach generates organelle signatures from millions of organelles across thousands of cells while preserving the native spatial context. Across metabolically specialized organs, organelle architecture distinguished major cellular identities, indicating that intracellular organization alone provides an informative and scalable readout of cell state.

The liver provided an ideal model system in which to test whether organelle architecture could resolve functional diversity within a single cell type. Hepatocytes are exposed to well-established gradients of oxygen, nutrients, and hormones along the portal vein-to-central vein axis, and classical models of liver zonation have emphasized a layered spatial partitioning of metabolic functions across this axis. Based on this, many analytical frameworks summarize zonation as averaged molecular profiles across ordered spatial bins, an approach that captures positional trends but can compress differences among hepatocytes within the same region (*35–37*, *48*, *49*). Our results refine this view by showing that, within the canonical liver zones, hepatocyte states with different organelle architectures coexist within local regions of the lobule. Rather than forming locally homogeneous layers, distinct hepatocytes spatially intermix forming a pattern of sub-zonal functional diversity, revealing a previously underappreciated degree of local heterogeneity across the hepatic acinus. This pattern suggests that hepatocyte function is organized not only by position along a continuous zonal gradient, but also by variation in intracellular architecture among neighboring cells that together coordinate metabolic tasks. Such coordination could enhance tissue plasticity by allowing the liver to maintain physiological functions while increasing tolerance to acute and chronic environmental stress (*16*, *85–87*).

Integration with spatial proteomic data further supported this idea. Cross-modal alignment linked organelle architectures to inferred proteomic programs associated with mitochondrial, peroxisomal, and lipid metabolic functions. The strong relationships observed support the interpretation that organelle-defined hepatocyte states correspond to complementary metabolic programs. More broadly, these analyses suggest that distinct neighboring hepatocytes within the same zone may contribute complementary metabolic activities, thereby distributing functional labor locally rather than acting as uniform zonal units. Together, these data are consistent with a model in which sub-zonal diversity potentially supports efficient division of metabolic labor within the liver. Considering this model, the follow up question arises: How is this sub-zonal diversity established? In addition to PV–CV gradients of oxygen, hormones, and nutrients, local cues related to bile canalicular organization, hepatocyte polarity, and interactions with Kupffer cells or neuronal inputs may shape organelle-defined states (*88–90*). Future studies combining *sOrganellomics* with targeted staining of these structures and other cell types should help resolve these contributions.

Applying *sOrganellomics* to nutritional perturbations showed that this sub-zonal hepatocyte spatial organization is plastic. Acute fasting induced coordinated remodeling of mitochondria, peroxisomes, and lipid droplets and generated distinct fasting-associated hepatocyte states across the lobule. Intravital imaging further linked these structural changes to increased mitochondrial membrane potential *in vivo*. These responses were disproportionately stronger near the central vein, reducing the PV-CV bioenergetic gradient while partially preserving sub-zonal diversity. This regional bias was consistent with the organelle-based transition landscape inferred from fixed tissue imaging. In contrast, chronic WD feeding reduced hepatocyte diversity and compressed the sub-zonal organization, consistent with a shift toward more uniform cellular states. These findings indicate that short-term and chronic metabolic stress differentially reshape intracellular architecture, with unique consequences for the cell state diversity of tissues. Together, these observations support the view that remodeling of subcellular architecture and local cell state diversity can serve as a sensitive readout of tissue status under changing physiological conditions.

Although some of these adaptations to nutritional perturbations may be reversible under acute conditions, sustained metabolic stress may reduce tissue resilience through loss of functional diversity. This possibility may be relevant to disease vulnerability, as WD-induced MASLD has been linked to hepatocellular carcinoma, and hepatocytes near the central vein—where we find diversity to be the lowest—are more susceptible to tumor initiation (*91*). These observations support a close relationship between a lower diversity of cell states and tissue susceptibility, that could potentially be captured by measurements of organelle architecture. Prior work supports this possibility, demonstrating that organelle remodeling *in vivo* can influence tissue function directly (*18*). Consistent with this, our predictive analyses further indicate that multi-organelle architecture contains sufficient information to classify hepatocyte states, nutritional condition, and early disease stages. This ability to classify early WD-associated changes also suggests that pathological remodeling can be detected before the most severe tissue alterations emerge. Whether organelle signatures can anticipate later functional decline or disease progression will require additional longitudinal and perturbational studies, but the present results highlight the potential of organelle architecture as a quantitative marker of tissue state.

More broadly, our findings highlight organelle architecture as complementary to molecular profiling approaches, providing a functional readout that is not fully captured by molecular measurements alone. Although transcriptomic methods have greatly advanced the study of cellular diversity, transcript abundance often correlates only modestly with protein levels, suggesting that transcript levels do not necessarily reflect functional cellular activity (*92*, *93*). Consistent with this distinction, proteomic heterogeneity patterns closely mirrored those derived from organelle architecture, suggesting that both modalities capture related aspects of the same underlying cellular states. At the same time, organelle-based analysis resolved finer transitions between heterogeneity domains potentially reflecting the greater cellular sampling achieved by *sOrganellomics*. Benchmarking of organelle-derived signatures with published spatial transcriptomic and proteomic datasets further supported the robustness of this approach (Fig. 2I) (*57*, *58*). A limitation of the current workflow is that the cell states it can resolve depend on the quality of the images and segmentation performance. However, *sOrganellomics* can be applied to other organelles and is readily adaptable across tissues and experimental conditions provided they can be imaged and segmented (see Methods).

Altogether, our study establishes organelle architecture as a measurable dimension of cellular state and introduces *sOrganellomics* as a scalable workflow for mapping functional diversity *in situ*. By combining high-resolution imaging with automated analysis, this approach enables systematic assessment of organ adaptation across biological scales—from organelle organization to tissue architecture—while preserving spatial context. Because *sOrganellomics* is compatible with conventional imaging platforms such as confocal microscopy and can, in principle, be extended to additional modalities, including volumetric electron microscopy and expansion microscopy, it provides a flexible and cost-effective strategy for interrogating functional diversity of cells across biological systems. Beyond its immediate applications, this workflow will facilitate generation of large multi-scale datasets that can be leveraged for data integration, mechanistic modeling, and machine learning-based prediction of cellular and tissue behavior. As imaging and molecular profiling continue to converge, systematic mapping of multi-organelle architecture may help bridge molecular regulation and cellular physiology, providing new insight into how intracellular structure contributes to tissue adaptation and disease.

## Acknowledgments

We would like to thank the Janelia Shared Resources for their excellent technical support. We are grateful to Meng Wang, Daniel Colón-Ramos, Wyatt Korff and Ron Vale for their feedback during the development of this project. We thank Mark Aronson from the Sgro Lab for his assistance in writing the script to crop acini from the liver lobule, Triveni Menon for revision of this manuscript, Joaquin Lilao-Garzón for critical feedback about the experiments, and Cedric Allier from the Saalfeld Lab for great discussions. We thank Alison Howard and Victoria Custard for their administrative support. The illustrations in Fig1. A and fig. S1 were created with <u>Biorender.com</u>.

## Author contributions

Conceptualization: D.F, methodology: D.F., A.H., R.A., investigation: D.F., R.A., A.H., A.D.J., machine learning implementation and analysis: D.F., R.A., A.H., benchmarking of spatial omics: D.F., S.M.G., writing, review, and editing: D.F., I.E.-M and R.A., visualization: D.F., R.A., supervision: D.F., J.F., project administration: D.F., J.F., funding acquisition: D.F., J.F.

## Data and material availability

All data are available in the main text or the supplementary material. Analysis code described in Materials and Methods section is available on GitHub (https://github.com/ahillsley/liv_zones, https://github.com/funkelab/motile).

## Materials and Methods

### Animals and Experimental Design

Eight-week-old Pham-excised (strain #018397) mice carrying the Mito-Dendra2 transgene (heterozygous males and females) were obtained from Jackson Laboratories. Breeding between heterozygous males and females was performed to obtain homozygous males. Homozygous males were then crossed with C57BL/6J wild-type females to generate heterozygous progeny for experimental use. Breeding was conducted in the vivarium at Janelia Research Campus. Only male mice were used to minimize variability. All procedures followed NIH guidelines and were approved by the Institutional Animal Care and Use Committee (Protocol # 25-0280) at Janelia Research Campus, Howard Hughes Medical Institute. Mice were housed under a 12-hour light/dark cycle with access to food and water ad libitum. All the livers were harvested at ∼ 6 am.

For nutritional perturbation experiments, animals were maintained on a control diet (TestDiets 58B0) until 6 weeks of age. Control mice (n=4) were continued on this diet until 10 weeks of age. Overnight-fasted mice (n=3) were deprived of food at 6 pm the evening prior to sacrifice and harvested at 6 am the following morning. Western diet (WD) mice (n=3) were switched to a high-fat diet (TestDiets 5GDD) at 6 weeks of age and maintained on this diet until 10 weeks. All the livers were harvested at ∼ 6 am.

For MASLD progression experiments, mice were placed on the high-fat diet starting at 6 weeks of age, and liver samples were harvested at 0, 7, 17, 31, 42, and 50 days of feeding (n=2 mice for each time point) retrospectively to match the final age. All the livers were harvested at ∼ noon.

Organs (liver and pancreas) were harvested after cardiac perfusion with 1x PBS to clear blood, followed by perfusion with 30 ml of 4% paraformaldehyde (PFA) at a controlled rate of 2.5 ml/min to prevent endothelial disruption. Harvested livers and pancreas were immersed in 4% PFA for 24 hours, washed three times with 1x PBS, and stored in 1% PFA until immunostaining.

For intravital microscopy, 10-weeks-old mice (n=3 for each group) were injected via lateral tail vein with 250 μL of 200 μM BODIPY 665/676 (Thermo Fisher Scientific Cat no: B3932) 24 hours before imaging to label lipid droplets. Retroorbital injection of 100 μL of 200 μM Tetramethylrhodamine, methyl ester (TMRM) (Thermo Fisher Scientific Cat no: T668) was performed just before the imaging session to measure mitochondria membrane potential changes. Food was removed at 6 pm the night before imaging for the fasted groups. Imaging was done next day at 8 am ±2 hours.

### Tissue Preparation and Immunostaining

Liver and pancreas samples were embedded in 4.6% and 3% low–melting-point agarose, respectively, and sectioned into 120 µm slices using a Leica vibratome (Leica VT1200S). Sections were permeabilized and blocked in PBS containing 0.5% Triton X-100 and either 10% fetal bovine serum (liver) or 2% bovine serum albumin (pancreas) for 1–2 h at room temperature. Tissue sections were incubated with primary antibodies for 48 h (liver) or 72 h (pancreas) at 4 °C in blocking buffer with gentle agitation. For liver samples, peroxisomes were labeled using mouse anti-PMP70 (MilliporeSigma, SAB4200181; 1:75). Additionally, rabbit anti-aldolase B (Invitrogen PA5-30218; 1:100) and mouse anti-CYP2E1 67263-1-Ig; 1:100) antibodies were used for liver to confirm metabolic zonation. For pancreas samples, primary antibodies included chicken anti-GFP (Aves Labs, GFP-1020; 1:1000) to enhance mito-Dendra2 fluorescence, rabbit anti-PEX14 (Proteintech 10594-1-AP; 1:50) for peroxisomes, guinea pig anti-glucagon (LSBio LS-C202759-100; 1:1000) to label pancreatic α cells, and rat anti-insulin (R&D Systems MAB1417; 1:500) to label pancreatic β cells. Sections were washed three times in PBS (10 min each) following primary antibody incubation. Secondary antibodies were applied overnight at 4°C. The following secondary antibodies were used (all 1:500): anti-mouse Alexa Fluor 647 (Thermo Fisher A21235), anti-chicken Alexa Fluor 488 (Invitrogen A78948), anti-rabbit Alexa Fluor 647 (Invitrogen A-31573), anti-guinea pig Alexa Fluor 790 (Jackson ImmunoResearch 706-655-148), and anti-rat Alexa Fluor 405 (Invitrogen A48268). Cell boundaries were labeled using Alexa Fluor 555 phalloidin (Thermo Fisher A30106; 1:100), and lipid droplets were visualized using HCS LipidTOX Red (Thermo Fisher H34476; 1:100). After secondary incubation, sections were washed in PBS and briefly post-fixed (pancreas only) in 4% PFA. For optical clearing, sections were incubated in EasyIndex (LifeCanvas Technologies, EI-500-1.52). Liver sections were incubated in 50% EasyIndex for 1 h followed by 100% EasyIndex for 3–5 h, whereas pancreas sections were incubated in 100% EasyIndex for 1–2 h. LipidTOX Red (1:100) was maintained during the clearing steps. Cleared sections were mounted on glass slides using Secure-Seal spacers (Thermo Fisher, 0523073) and imaged immediately.

### Confocal Imaging

Images were acquired using a Leica Stellaris 8 confocal microscope. For each liver section, at least three lobules were selected and three acini per lobule spanning the portal vein (PV) to central vein (CV) axis were imaged. Portal and central vein regions were identified based on vascular morphology and confirmed using phalloidin staining to visualize hepatocyte boundaries. For the pancreas, images were acquired from control mice (n=3) including both endocrine (confirmed by glucagon and insulin expression) and exocrine regions. At least 25 such regions among 3 animals were imaged for analysis. Z-stacks covering approximately 10 μm in depth (∼50 optical sections) were acquired for each region. Imaging was performed using a 63× oil-immersion objective, with a pinhole size of 0.5 Airy units, a zoom factor of 2, and an image resolution of 2048 × 2048 pixels.

### Intravital Microscopy

For intravital imaging, mice were anesthetized and a small surgical incision was made to expose the lateral liver lobe. The exposed liver surface was gently positioned against a glass coverslip mounted on the microscope stage to stabilize the imaging plane. A Secure-Seal spacer (Thermo Fisher Scientific, Cat. No. S24737) containing a single well was placed on the coverslip, and the imaging region of the liver was positioned within the well. The edges of the liver tissue were secured to the spacer using surgical adhesive (bio-glue) to minimize motion during acquisition. Imaging was performed using a 40× oil-immersion objective with a pinhole size of 0.75 Airy units, a zoom factor of 2, and an image resolution of 2048 × 2048 pixels. Regions of the liver with active blood flow were used to detect PV and CV region and regions around PV and CV were imaged. Animals remained under anesthesia throughout the imaging session and were euthanized immediately after completion of image acquisition.

### Image preprocessing, organelle Segmentation and Feature Extraction (sOrganellomics workflow)

Images were processed using the Liv_zones pipeline. PV and CV coordinates were manually selected for every lobule in the livers. Acinar regions spanning from PV to CV were cropped from each lobule. Each resulting acinus was normalized to a unit length (−1 to1), and hepatocyte centroids were projected onto this axis based on their relative position between the PV (1) and CV (−1). For zonation analyses, this normalized axis was partitioned into three equal segments corresponding to zone 1 (periportal), zone 2 (midzonal), and zone 3 (pericentral). Non-parenchymal liver cell types were excluded based on cell size and mitochondrial content to focus on hepatocyte state diversity. For pancreas, endocrine regions were confirmed by insulin and glucagon staining, while the exocrine regions were identified morphologically in phalloidin-stained pancreatic sections. Segmentation was performed using Cellpose v2 with separate custom models for pancreas and liver. Models were trained on top of cellpose cyto2 model and custom models were generated each for mitochondria, peroxisomes, and lipid droplets using supervised human-in-the-loop annotation on representative image crops. Segmentation accuracy among organelles ranged from ∼72% to 96% based on comparison to manual annotations.

Organelle features were quantified from organelle masks using skimage.measure.regionprops_table in scikit-image. Extracted morphological features included area, perimeter, centroid coordinates, major and minor axis length, and solidity. Derived metrics included aspect ratio, circularity, organelle-to-cell assignment, distance to the hepatocyte boundary, and downstream cell-level summaries such as density. Due to the wide dynamic range of lipid droplet sizes, segmentation was performed with three diameter parameters (16, 132, and 645 pixels) to capture small, medium, and large droplets. An additional model of peroxisome was generated to account for large, clustered peroxisomes in the pancreas that are not distinguishable at single organelle level and named them as large peroxisomes. Features were aggregated at the single-cell level by averaging across z-stacks. Cells were linked across slices using Motile (https://github.com/funkelab/motile).

### Clustering and Cell State Identification

Single-cell organelle feature matrices were standardized using z-score normalization. Principal Component Analysis (PCA) was performed to capture the major axes of variation and to enable interpretable feature contributions underlying cluster structure. Clustering was then performed using Gaussian mixture models (GMMs) on the PCA embeddings. To determine the appropriate number of principal components (PCs) for clustering, we evaluated cluster stability across varying numbers of PCs using the mean Adjusted Rand Index (ARI) across bootstrap iterations. The first two PCs yielded the highest stability and were therefore selected for downstream clustering. To avoid overfitting or underfitting in cluster assignment, the optimal number of clusters was determined using a multi-metric GMM approach combining elbow method, Bayesian information criterion (BIC), Akaike information criterion (AIC), silhouette score, and cross-validation. PCA embeddings were additionally used for visualization.

### Benchmarking of spatial omics modalities

Spatial Organellomics was benchmarked against spatial transcriptomics and spatial proteomics using a standardized computational framework that evaluates clustering robustness, separation, and resolution under matched features and cell budgets. All preprocessing and analyses were implemented in Python using Scanpy and custom scripts.

Transcriptomic data were normalized to 10,000 counts per cell, log-transformed, restricted to the top 2000 highly variable genes, and scaled. Proteomic data were median-normalized across proteins, log₂-transformed, and mean-centered. Organelle-derived features from *sOrganellomics* were scaled across cells. To ensure comparability across modalities, features were ranked by variance calculated prior to scaling and the top 98 features were retained for each dataset, corresponding to the smallest post-processing feature set across modalities (*1*). Datasets were further subsampled to 390 cells, matching the smallest dataset size and eliminating sample-size– driven differences in clustering performance (*2*, *3*). PCA was performed following feature selection.

Clustering was performed using two independent strategies to reduce algorithm-specific bias. Leiden clustering was applied to a k-nearest-neighbor graph constructed in PCA space using cosine distance for transcriptomics and Euclidean distance for proteomics and *sOrganellomics* (resolution = 0.5). In parallel, multi-metric GMM was performed as described earlier (*4–8*).

Clustering robustness was assessed by bootstrap resampling. GMM models were refit on random subsamples of the data and robustness was quantified as the ARI between bootstrap-derived assignments and those obtained from the full dataset. Cluster separation was evaluated using maximal pairwise overlap in GMM posterior responsibilities and the minimum Mahalanobis distance between component means. Assignment certainty was calculated as one minus the normalized entropy of posterior probabilities.

Clustering resolution was quantified using silhouette score and Davies–Bouldin index, both scaled to bounded values between 0 and 1. An additional effective resolution metric combined cluster-size granularity (effective number of clusters derived from the cluster size distribution) with stability across neighboring component numbers, measured as the ARI between solutions obtained at *k* ± 1.

All metrics were scaled to the range [0,1] and combined into a composite clustering score using fixed weights: robustness (0.15), overlap-based separation (0.15), Mahalanobis separation (0.15), silhouette-based resolution (0.15), Davies–Bouldin–based resolution (0.15), assignment certainty (0.15), and effective resolution (0.10) (*9*). Composite score uncertainty was estimated by bootstrap resampling, recalculating clustering metrics for each iteration and reporting the mean and standard deviation across bootstrap replicates for each modality.

### Zonation model validation

To evaluate whether hepatocyte identity is fully explained by spatial position along the PV–CV axis, we quantified multiple per-acinus metrics assessing positional determinism of hepatocyte categories (CAT). These analyses measure (i) whether residual spatial structure remains after accounting for position, (ii) the predictive sufficiency of position alone, (iii) the fraction of category variability explained by position, and (iv) whether local cell–cell organization exceeds expectations from zonation. All analyses were restricted to control livers.

Spatial position was represented by a continuous normalized PV–CV coordinate for each hepatocyte and discretized into *n* equally spaced spatial bins (default *n* = 30). Metrics were computed independently for each acinus to avoid pooling artifacts across lobules.

Residual spatial autocorrelation was first quantified using Moran’s I applied to hepatocyte category residuals. For each cell and category, residuals were defined as *r*_*i*_ = *y*_*i*_ − *P*^)^(CAT ∣ *position*),where *y*_*i*_denotes the observed indicator for a given hepatocyte category and *P*^)^(CAT ∣ *position*) represents the expected category probability within the corresponding spatial bin. Spatial neighbors were defined using a k-nearest-neighbor graph (*k* = 8). Moran’s I values were calculated on residuals and averaged across hepatocyte categories. To test significance, hepatocyte category labels were permuted within spatial bins (200 permutations per acinus), preserving zonation structure while disrupting local spatial organization. Residual Moran’s I values were compared with this null distribution using two-sided tests. Significant residual Moran’s I indicates spatial clustering beyond that expected from positional zonation alone.

Next, we quantified the predictive sufficiency of spatial position using a Bayes-optimal classification accuracy metric. For each spatial bin, the most frequent hepatocyte category was assigned as the predicted label. Overall classification accuracy was calculated as

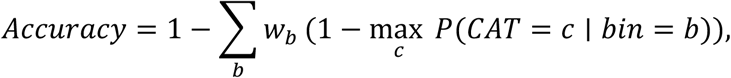

where *w*_*b*_represents the fraction of cells in spatial bin *b*. This metric represents the theoretical maximum classification performance achievable using spatial position alone.

We then quantified the fraction of hepatocyte identity explained by spatial position using conditional entropy. For each acinus,

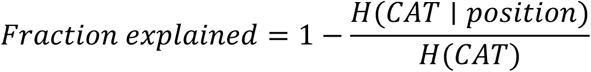

where *H*(CAT)is the Shannon entropy of hepatocyte categories within the acinus and *H*(CAT ∣ *position*)is the entropy conditional on spatial bin. Values range from 0 (identity independent of position) to 1 (identity fully determined by position). Statistical significance was assessed using within-acinus permutation testing (500 permutations) in which category labels were shuffled while preserving spatial coordinates. Two-sided permutation p-values were calculated as

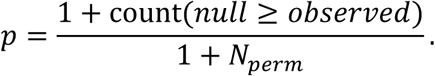

The unexplained component of identity variability was defined as 100 x (1-Fraction explained).

To assess spatial organization beyond zonation, we quantified position-conditioned neighbor enrichment. For each pair of hepatocyte categories, observed adjacency counts were compared with a null distribution generated by permuting category labels within spatial bins (200 permutations per acinus). Significant enrichments (p < 0.05) were counted per acinus. Under strict positional determinism, local adjacency structure should be fully explained by shared spatial bins; excess significant neighbor interactions indicate organization beyond positional zonation.

Group-level summaries were obtained by averaging metrics across acini. Confidence intervals were estimated using bootstrap resampling across acini (2000 replicates). For visualization in fig S13, selected metrics were linearly transformed to zero-centered scales to facilitate comparison across conceptually distinct measurements. These transformations were applied after statistical testing and therefore do not influence permutation tests, bootstrap confidence intervals, or Bayesian inference.

Normalization anchors were chosen to provide interpretable reference points. For the fraction-explained metric, zero corresponds to a reference value of 70% positional determinism. For position-only classification accuracy, zero corresponds to an 80% predictive accuracy threshold.

For residual Moran’s I, normalized values represent the log10 ratio between the permutation p-value and the conventional significance threshold (0.05), such that negative values indicate statistically significant residual spatial structure. For neighbor enrichment, zero corresponds to the expected number of significant neighbor interactions under the null hypothesis (5% false positives), with negative values indicating excess local organization beyond zonation expectations.

### Heterogeneity Metrics

Simpson’s dominance index and Shannon entropy were calculated per spatial bin based on category proportions. For *sOrganellomics* data, metrics were computed per acinus and summarized across acini. For spatial proteomics data, metrics were computed from proteomics-defined clusters using identical spatial binning.

### Cross-modal proteomic alignment and inference

To infer proteomic programs associated with *sOrganellomics* states, we integrated independently generated spatial proteomics data with *sOrganellomics* measurements collected from different animals and experiments. Because no shared cells and no shared acini were available across modalities, alignment was performed at the level of discrete hepatocyte states (categories) rather than at the single-cell level. Cells in each modality were clustered into five spatial categories (*sOrganellomics*: OH1–OH5; proteomics: PH1–PH5) and each cell was assigned an acinus coordinate along the portal–central axis.

To make acinus coordinates comparable across experiments, the position coordinate *x* was normalized within each acinus by min–max scaling to [0,1] (with the raw coordinate retained for reference). For each OH–PH category pair, spatial correspondence was quantified using the first Wasserstein distance (1D optimal transport) between their category-specific *x*-distributions (*10*, *11*). Because acini are experiment-specific and not matched across modalities, OH–PH distances were computed by averaging Wasserstein distances over the cross-product of acini pairs (optionally subsampling acinus-pairs per OH–PH entry for computational efficiency, recorded as support counts). This yielded a 5 × 5 distance matrix *W*(rows OH, columns PH). For descriptive purposes, an optimal one-to-one OH–PH matching was obtained by the Hungarian algorithm applied to *W* (*12*). Statistical significance of the observed matching cost was assessed using an acinus-preserving permutation test (1,000 permutations), in which OH category labels were randomly permuted within each acinus, followed by recalculation of the distance matrix and Hungarian cost to generate a null distribution (*13*).

Because the only quantity shared across modalities is spatial position, the final soft correspondence was derived from spatial evidence only (to avoid circularity from proteomic features). The Wasserstein distance matrix *W*was converted to a soft spatial support matrix by applying an exponential kernel,

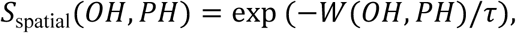

where *τ*(kernel temperature) was set to the median of all entries in *W*. Each OH row was then row-normalized to sum to 1. Two additional, independent spatial supports were computed: (i) rank support, based on agreement between OH and PH category ordering along the axis using median *x*(support ∝ 1/(1+∣ rank(*OH*) − rank(*PH*) ∣)); and (ii) interval overlap support, computed from the overlap of the 10–90% quantile intervals of category-specific *x*-distributions (overlap/union). Each support matrix was row-normalized, and the three supports were combined into a CONSENSUS OH→PH correspondence matrix by a weighted average, followed by final row normalization. CONSENSUS entries can be interpreted as the probability of PH conditioned on OH.

Proteome inference was performed by computing PH category mean protein abundance profiles from spatial proteomics and projecting them onto OH categories using CONSENSUS weights. To reduce batch effects in the proteomics dataset, protein abundances were corrected using a batch-aware mean-shift procedure (per protein: subtract batch mean and add the global mean) before computing category means. The inferred proteome for each OH was then computed as a weighted sum of PH means:

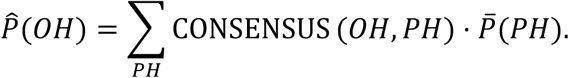

Downstream pathway scores were computed from inferred OH proteomes as the mean abundance of proteins in predefined gene sets and visualized either as relative contributions (within-pathway normalization) or as z-scored enrichment across OH categories.

Alignment quality and robustness were evaluated using complementary diagnostics: (i) distribution reconstruction error, quantifying how well each OH *x*-distribution is explained by the PH mixture implied by CONSENSUS; (ii) ordering agreement, comparing OH median *x* to the CONSENSUS-weighted expected PH median *x* (Spearman rank correlation); and (iii) CONSENSUS sharpness, summarized by normalized row entropy (*H* = −∑*p*log *p*/log *K*) (*14*). Uncertainty was quantified by bootstrap resampling of cells within categories (recomputing *W*, *τ*, CONSENSUS, and inferred proteomes per resample) to obtain confidence intervals and stability metrics (*15*), including bootstrap frequencies of hard OH–PH matches and correlations between bootstrap and baseline CONSENSUS rows. Sensitivity analyses were performed by varying *τ*(e.g., 0.5×, 1×, 2×) and by varying the relative weights of the three support components, reporting stability of the resulting CONSENSUS mappings.

### Nitrogen Functional Indices

To quantify nitrogen handling specialization across *sOrganellomics*-defined hepatocyte states, we derived composite functional indices from the inferred OH proteomes obtained by the cross-modal inference framework described above. Nitrogen-related metabolic modules were defined using curated protein sets representing major components of hepatic nitrogen metabolism, including amino acid degradation programs, ammonia-generating reactions, urea-cycle enzymes, glutamine synthetase activity, and one-carbon–associated modules. For each module (or sub-pathway), a pathway score was computed as the mean abundance of its constituent proteins within each inferred OH proteome.

To enable combination of distinct pathways on a comparable scale, pathway scores were standardized by z-scoring across OH categories (performed independently for each pathway). Composite indices were then computed as linear combinations of these z-scored pathway scores to represent relative functional balance between nitrogen-generating and nitrogen-disposal processes. Specifically, an aggregate amino acid breakdown program score (AA_breakdown) and an aggregate ammonia-generation program score (Ammonia_gen) were computed by averaging z-scored pathway values across the relevant sub-pathways. Urea-cycle activity (Urea_cycle) and glutamine synthesis activity (Glutamine_syn; GLUL module) were similarly defined.

Using these standardized program scores, the indices were defined as:

- **Net Amino Acid Catabolism:**

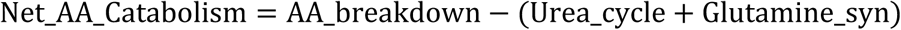

- **Ammonia Release Index** (net ammonia pressure after detoxification pathways):

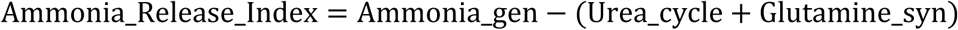

- **Urea Release Bias** (preference for urea-cycle disposal vs glutamine synthesis):

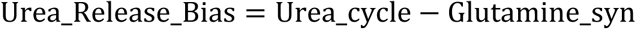

- **Glutamine Release Bias** (preference for glutamine synthesis vs urea-cycle disposal):

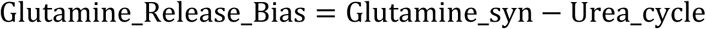

These indices represent inferred functional bias from protein abundance patterns rather than direct metabolic flux. They were used to rank and classify hepatocyte states according to dominant nitrogen-handling tendencies and were visualized across OH states using heatmaps and dot/bubble plots (Figure 3K).

### Trajectory Analysis

Single-cell organelle feature matrices from control and fasting hepatocytes were combined after z-score normalization across cells. Principal component analysis (PCA) was performed on the merged dataset to obtain a low-dimensional representation of the integrated organelle feature space. A probabilistic transition model was constructed using a forward Markov framework by first generating a k-nearest-neighbor graph in PCA space based on Euclidean distances and then converting edge weights into transition probabilities through row normalization of the adjacency matrix. Pseudotime (Shown in Figure 5B as Transition score) values were inferred from the resulting transition matrix using iterative propagation of transition probabilities across the graph, yielding a continuous ordering of hepatocytes along the inferred remodeling trajectory. The dominant remodeling axis was defined by the first principal component (PC1). Association between trajectory progression and structural remodeling was quantified by Spearman rank correlation between transition probability and PC1 projection. For spatial analysis, hepatocyte PC1 projections were grouped into normalized portal vein–to–central vein (PV–CV) spatial bins derived from acinus alignment, and mean PC1 values were computed per bin and summarized across replicates, with variability reported as ± SEM.

### Intravital imaging analysis and structure–function coupling

Intravital microscopy datasets were processed to extract single-cell mitochondrial and lipid droplet (LD) features and quantify their relationship with mitochondrial membrane potential (TMRM). Hepatocytes were segmented using a semi-automated workflow in Napari (v2), and organelle segmentation and feature extraction were performed using the Liv-Zones package. In addition to organelle features mitochondrial membrane potential was also quantified as normalized TMRM intensity (TMRM/Dendra2 fluorescence).

TMRM-defined bioenergetic states were defined by fitting Gaussian mixture models to log₁₀-transformed normalized TMRM intensities from control hepatocytes, and the resulting thresholds were applied to both control and fasting datasets.

For structure–function coupling analysis, mitochondrial and LD features were standardized (z-scored) across all hepatocytes. Feature weights were derived from the absolute Pearson correlation between each feature and TMRM intensity, normalized within each organelle group, with directionality determined by the sign of the correlation. Weighted mitochondrial and LD axes were computed as the sum of signed, weighted standardized features. A global organelle architecture score was generated by combining these axes using coefficients from linear regression of TMRM intensity across all cells.

Structure–function coupling was evaluated by regressing normalized TMRM intensity against the global organelle score separately for periportal (PV) and pericentral (CV) hepatocytes using ordinary least squares regression. Pearson correlation coefficients, slopes, and coefficients of determination (R²) were reported with 95% confidence intervals. Robustness was assessed using five-fold cross-validation with the global structural geometry fixed and regression parameters re-estimated in each fold.

### Machine Learning

Multilayer perceptron (MLP), Random Forest, and multinomial Logistic Regression models were trained using organelle-derived morphological features as inputs. All features were normalized to a 0–1 range using min–max scaling, and the dataset was randomly split into training (80%) and validation (20%) sets. A total of 10,542 hepatocytes across three nutritional regimens and ∼21,000 hepatocytes across Metabolic Dysfunction-Associated Steatotic Liver Disease (MASLD) progression were included in the analysis. Separate MLP models, consisting of three hidden layers with 64 neurons each, were trained to predict hepatocyte category, nutritional regimen, and MASLD progression stage. Random Forest and Logistic Regression models were trained to predict hepatocyte category and nutritional regimen using an ensemble of decision trees and a multinomial formulation, respectively. Model performance was evaluated on the validation set using overall accuracy and row-normalized confusion matrices.

## Code Availability

The Liv-Zones pipeline and trained segmentation models are available at https://github.com/ahillsley/liv_zones. The Motile package for cell linking is available at https://github.com/funkelab/motile.

