## Supplementary Figures and Legends for "Spatial Organellomics Maps Cell State Diversity and Metabolic Adaptation in Tissues"

**Affiliations:**

**Table of contents**

**Supplementary figures and legends**

**Fig. S1. Schematic representation of *sOrganellomics* pipeline**. **A,** The pipeline uses immunolabeled volumetric images of organelles and cell boundaries obtained from tissue sections. Pancreatic sections are additionally immunostained for glucagon and insulin to identify endocrine cells in the Islets of Langerhans (region marked by pink dashed lines in the glucagon and insulin panels). EN: endocrine Islet, EX: exocrine cells. For liver sections, the X and Y coordinates of the portal vein (PV) and central vein (CV) are manually obtained. These coordinates are then used in Python to crop large images into individual acini, each comprising a single PV and CV. **B,** Custom segmentation models are developed by starting with the built-in Cellpose cyto model and optimizing it using a human-in-the-loop approach. These models are applied to obtain cell and organelle masks across tissues, achieving high-precision segmentation. **C,** The refined models, packaged as *Liv_Zones*, are called via Python, where the processed images are used as input to generate masks for each cell within a tissue and for the organelles of interest within each cell. These masks are used to extract features listed in the table. Feature extraction produces CSV files either for each organelle separately or as averaged properties at the cell level. All masks retain positional information relative to the PV and CV within the tissue. These CSV files are subsequently used to visualize the distribution of organelle features, analyze spatial trends across the tissue, and classify cells based on organelle characteristics.

**Fig. S2. Differential organelle features among hepatocytes in the liver. A,** Hepatocyte masks color-coded for organelle densities. Normalized z-scores (color scale: white for -1 (lower values) to purple for +1 (higher values). **B,** Heatmap showing spatial organelle features distribution in the acini. The x-axis represents relative cell positions in the PV-to-CV axis, while the y-axis lists individual organelle features. Normalized z-scores (color scale: white for -1 to purple for +1) reveal spatial variations in organelle architecture, emphasizing the heterogeneity of subcellular structures across the acini. **C,** Volcano plots illustrating differences in organelle features between three functional zones in the liver. Each point represents a single organelle feature, with the x-axis showing the log2 fold change (logFC) between regions and the y-axis displaying the -log10 of the False Discovery Rate (FDR). Organelles with significant differences (FDR $< 0.05$) are shown as larger circles, with those enriched in one region colored purple (Z2, left, Z3, right) and those enriched in the other region colored blue (Z1). The pie charts above the plots summarize the contribution of the three measured organelles to the primary differences between the indicated liver zones (Z1, Z2, Z3).

**Fig. S3. Correlation matrix of key features between organelles.** Sketch of hepatic acinus (top) representing three functional zones (zone 1, zone 2 and zone 3). Each block corresponds to one liver zone, and within each block the matrices show pairwise correlations of organelle features for the indicated organelle pairs. Values close to 1 or -1 indicate strong positive or negative correlations, respectively, while values close to 0 indicate weak or no correlation.

**Fig. S4. Trend lines of average organelle features from PV to CV under control diet.** The distance from PV to CV was subdivided into 20 bins, and organelle features were averaged at each bin. Individual features were plotted separately as trend lines for mitochondria (top panel), peroxisome (middle panel), and lipid droplets (bottom panel) along the PV-CV axis. The x-axis represents the distance from the PV to CV, while the y-axis represents the average values of the organelle features. The shadow along the trend lines indicates the standard error of the mean (SEM).

**Fig. S5. Trend lines of average feature values for mitochondria subpopulations from PV to CV.** The mitochondria from control livers were subdivided into 3 subpopulations based on aspect ratio, and their features were plotted as trend lines from PV to CV into 20 bins. The x-axis represents the distance from the PV to the CV, while the y-axis represents the average values of the organelle features. The shadow along the trend lines indicates the standard error of the mean (SEM).

**Fig. S6. Trend lines of average feature values for peroxisome subpopulations from PV to CV axis.** The peroxisomes from control livers were subdivided into 3 subpopulations based on aspect ratio and their features were plotted as trend lines from PV to CV into 20 bins. The x-axis represents the distance from the PV to the CV, while the y-axis represents the average values of the organelle features. The shadow along the trend lines indicates the SEM.

**Fig. S7. Trend lines of average feature values for lipid droplet subpopulations from PV to CV axis.** The Lipid droplets from control livers were subdivided into 4 subpopulations based on area and their features were plotted as trend lines from PV to CV into 20 bins. The x-axis represents the distance from the PV to the CV, while the y-axis represents the average values of the organelle features. The shadow along the trend lines indicates the SEM.

**Fig. S8. Hepatocyte classification based on organelle signatures. A,** Principal component analysis (PCA) depicting hepatocyte distribution based on average organelle features at the single-cell level, mapped onto Principal Component 1 (PC1) and Principal Component 2 (PC2). **B,** Bar graph showing the contribution of each principal component to the explained variance. **C,** Cluster stability across varying numbers of principal components (PCs). The x-axis shows PCs (2–10), and the y-axis shows mean Adjusted Rand Index (ARI). Bars represent the mean across 15 bootstrap iterations per PC number; error bars indicate ± SEM. **D,** Contribution of organelle features to principal components (PC1 and PC2). The x-axis shows individual organelle features, and the y-axis shows their contribution to each principal component. **E,** Schematic representation of the strategy for unbiased selection of optimal cluster number using a multi-metric Gaussian Mixture Model (GMM) approach. **F,** PCA plots showing hepatocyte categories in individual liver samples. **G,** Percentage mean differences between hepatocyte categories. The means of organelle features for distinct hepatocyte categories on the y-axis were compared against each other, as indicated. A one-way ANOVA was conducted to assess significant differences across all groups. Examples of statistically significant features for each organelle are shown. Pairwise $P$-values were calculated. N.S., non-significant ($P > 0.05$) and *, $P < 0.05$ and ***, $P < 0.0001$.

**Fig. S9. PCA plots illustrating the distribution of key mitochondria features across five hepatocyte categories.** Each row corresponds to a specific mitochondria feature. The first column displays the overall distribution of the indicated feature among the five clusters in the PCA plot, while the subsequent columns highlight the distribution within individual clusters by emphasizing the corresponding hepatocyte category. The color scale represents normalized z-scores, ranging from-1 (white, indicating lower-than-average values) to +1 (purple, indicating higher-than-average values), capturing variations in mitochondria features both among and within clusters.

**Fig. S10. PCA plots illustrating the distribution of key peroxisomal features across five hepatocyte categories.** Each row corresponds to a specific peroxisomal feature. The first column displays the overall distribution of the indicated feature among the five clusters in the PCA plot, while the subsequent columns highlight the distribution within individual clusters by emphasizing the corresponding hepatocyte category. The color scale represents normalized z-scores, ranging from-1 (white, indicating lower-than-average values) to +1 (purple, indicating higher-than-average values), capturing variations in peroxisomal features both among and within clusters.

**Fig. S11. PCA plots illustrating the distribution of key LD features across five hepatocyte categories.** Each row corresponds to a specific LD feature. The first column displays the overall distribution of the indicated feature among the five clusters in the PCA plot, while the subsequent columns highlight the distribution within individual clusters by emphasizing the corresponding hepatocyte category. The color scale represents normalized z-scores, ranging from -1 (white, indicating lower-than-average values) to +1 (purple, indicating higher-than-average values), capturing variations in LD features both among and within clusters.

**Fig. S12. Comparison of clustering performance and zonal composition of hepatocyte categories across modalities. A,** Summary table of clustering performance metrics and dataset harmonization across modalities and methods. The first three rows correspond to Leiden clustering and the remaining three to Gaussian Mixture Model (GMM) clustering. Within each method, rows from the top represent analysis for dataset from *sOrganellomics* (O), proteomics (P), and transcriptomics (T), respectively. **B,** Stacked bar plots showing the proportion of hepatocyte categories derived from *sOrganellomics* across the three liver zones (Z1-Z3). **C,** PCA visualization of hepatocyte categories defined using spatial proteomics data, illustrating separation into five distinct categories. **D,** Stacked bar plots showing the proportion of hepatocyte categories derived from proteomics across the three liver zones (Z1 -Z3).

**Fig. S13. Statistical testing of the classical model of liver zonation.** Per-acinus model validation metrics (n = 30 acini) assess whether spatial position alone explains hepatocyte identity. Metrics include residual Moran's I, maximum classification accuracy based on position, fraction of variability explained by position, and position-conditioned neighbor enrichment (see Methods). Each dot represents an individual acinus; shaded regions indicate group means with 95% bootstrap confidence intervals. Values $< 0$ indicate statistically significant spatial organization beyond that explained by position, whereas values $> 0$ indicate spatial organization consistent with the null expectation. A summary on the right lists the models tested and highlights that hepatocyte diversity is only partially explained by spatial position.

**Fig. S14. Validation and robustness of cross-modality alignment used for proteome inference.** **A,** Baseline OH→PH consensus correspondence matrix used to transfer proteomic profiles from spatial proteomics hepatocyte (PH) categories to sOrganellomics hepatocyte (OH) categories. Each OH row is row-normalized to sum to 1 and represents a soft assignment over PH categories. Consensus weights combine three spatial evidences: (i) spatial distribution similarity derived from Wasserstein distances between within-acinus normalized zonation coordinate ($x$) distributions, converted to a similarity kernel with temperature parameter $\tau$; (ii) median-order agreement (rank support) based on category median $x$; and (iii) interval overlap support based on overlap of category $x$quantile intervals (10–90%). **B,** Distribution reconstruction error evaluating how well each OH category observed $x$-distribution is explained by the PH mixture implied by Consensus. Bars show Wasserstein distance between the OH distribution and the reconstructed PH-mixture distribution (lower indicates better reconstruction). **C,** Zonation ordering agreement between modalities under the soft correspondence. Points show OH median $x$(x-axis) versus the expected PH median $x$predicted by Consensus (y-axis), computed as the Consensus-weighted average of PH median positions. Spearman rank correlation (ρ) and two-sided p-value are shown. **D,** Bootstrap stability of hard category matching. Heatmap shows the frequency with which each OH category is assigned to each PH category across bootstrap resampling runs (Hungarian assignment performed on bootstrap-estimated Wasserstein distance matrices). Values represent bootstrap frequencies (0–1). **E,** Sensitivity to kernel temperature $\tau$. Heatmap reports, for each OH row, the correlation between the baseline Consensus row and the Consensus row recomputed after scaling $\tau$(e.g., 0.5×, 1×, 2×). Higher correlation indicates robustness to the choice of kernel softness.

**Fig. S15. Trend lines of average organelle features from PV to CV under different nutritional conditions.** Each graph includes three trend lines corresponding to control, fasting, and western diet conditions. Individual features were plotted separately as trend lines fitted using linear regression for mitochondria (top panel), peroxisome (middle panel) and lipid droplets along the PV-CV axis. The x-axis represents the distance from the PV to the CV divided into 20 bins, while the y-axis represents the average values of the organelle features. The shadow along the trend lines indicates the SEM.

**Fig. S16. Trend lines of average features for mitochondria subpopulations from PV to CV under different nutritional conditions.** Mitochondria are subdivided into 3 subpopulations based on their aspect ratio and their features are plotted as trend lines from PV to CV into 20 bins. Each graph includes three trend lines corresponding to control, fasting, and western diet conditions. The x-axis represents the distance from the PV to the CV, while the y-axis represents the average values of the organelle features. The shadow along the trend lines indicates the SEM.

**Fig. S17. Trend lines of average feature values for peroxisome subpopulations from PV to CV under different nutritional conditions.** Peroxisomes are subdivided into 3 subpopulations based on their aspect ratio and their features are plotted as trend lines from PV to CV into 20 bins. Each graph includes three trend lines corresponding to control, fasting, and western diet conditions. The x-axis represents the distance from the PV to the CV, while the y-axis represents the average values of the organelle features. The shadow along the trend lines indicates the SEM.

**Fig. S18. Trend lines of average feature values for lipid droplet subpopulations from PV to CV under different nutritional conditions.** Lipid droplets are subdivided into 4 subpopulations based on their area and their features are plotted as trend lines from PV to CV into 20 bins. Each graph includes three trend lines corresponding to control, fasting, and western diet conditions. Subtype 4 was more abundant in fasting and western diet, while they were uncommon in control. The x-axis represents the distance from the PV to the CV, while the y-axis represents the average values of the organelle features. The shadow along the trend lines indicates the SEM.

**Fig. S19. Reduced hepatocyte heterogeneity under nutritional perturbations. A, C, E,** PCA plots depicting the spatial distribution of hepatocyte categories along the PV–CV axis for control, fasting, and WD groups, respectively. The individual plots on the right illustrate the spatial distribution of each indicated hepatocyte category along the PV–CV axis, as represented by the color bar. **B, D, F,** Graphs showing the proportion of hepatocyte categories in control, fasting, and WD groups, respectively. **G,** Heatmaps displaying unique mitochondrial, peroxisomal, and lipid droplet (LD) properties across hepatocyte categories for each nutritional group. Organelles were classified into subtypes based on aspect ratio (mitochondria and peroxisomes) or area (LDs). The y-axis represents key features corresponding to mitochondria, peroxisomes, and LDs, while the x-axis denotes hepatocyte categories within the three nutritional groups. The color scale represents normalized z-scores, ranging from -1 (black, indicating lower-than-average values) to +1 (white, indicating higher-than-average values), highlighting variations in organelle signatures across hepatocyte categories. **H,** Graphs displaying the percentage of organelle subtypes as a fraction of the total organelle population for each hepatocyte category in control, fasting, and WD groups.

**Fig. S20. Trajectory analysis of hepatocyte state transitions and organelle remodeling. A,** Direct and indirect transition probabilities from control to fasting states. Bars show Markov absorption probabilities for observed control-to-fasting state transitions, decomposed into direct transitions and paths via intermediate states (H1–H5), as indicated by color. **B,** Forward transition probabilities from control state H4 to fasting states (FH1–FH4) resolved by transition path. The y-axis lists distinct direct and multi-step paths and bars indicate the corresponding Markov absorption probabilities. **C,** Schematic of inferred transition trajectories from control to fasting states. Control hepatocyte states converge through a transition hub (H4 and FH2) and diverge toward downstream fasting states (FH1, FH3 and FH4). An example of organelle remodeling is shown for H1 to FH1 transition. **D,** Loadings of organelle features on the first principal component (PC1) representing the dominant remodeling axis. Bars indicate the contribution and direction of each feature to the PC1.

**Fig. S21. Classification of hepatocyte category and nutritional regimen using Logistic Regression and Random Forest models**. **A, B** Confusion matrices showing classification performance of multinomial Logistic Regression models for hepatocyte category (A) and nutritional regimen (B). **C, D** Confusion matrices showing classification performance of Random Forest models for hepatocyte category (C) and nutritional regimen (D). Values in each box represent row-normalized percentages, indicating the proportion of cells in each true class assigned to each predicted class. Rows correspond to true labels and columns to predicted labels.

Supplementary Figure 1

#### A. Image acquisition and processing

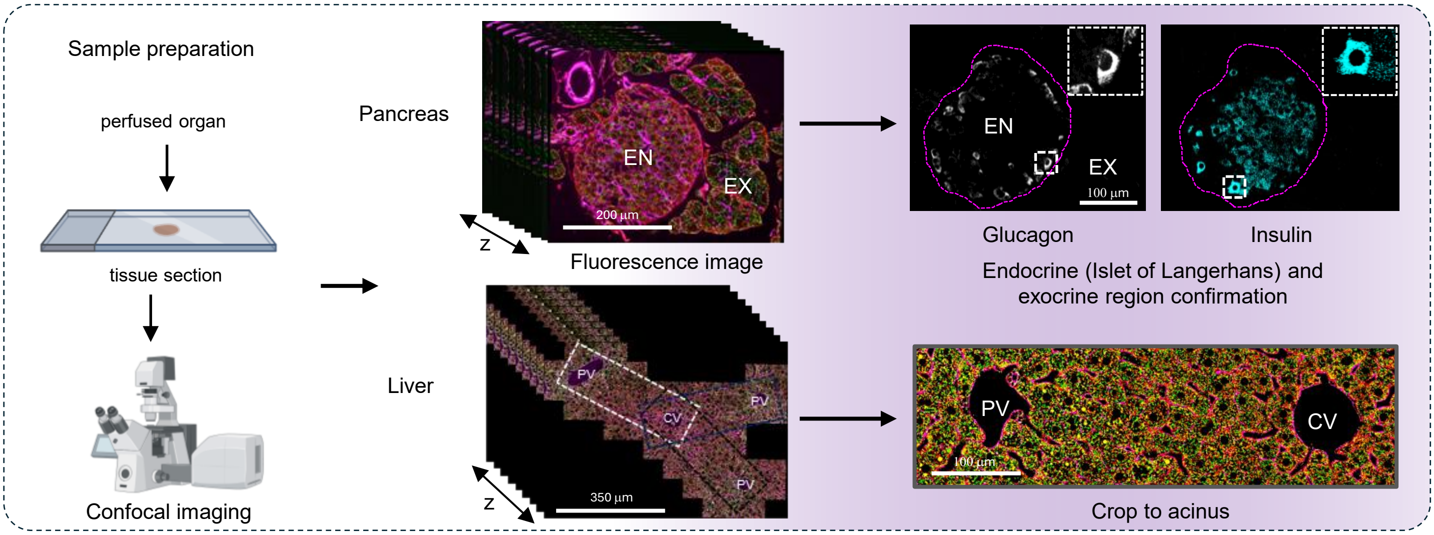


#### B. Cell and organelle segmentation models generation

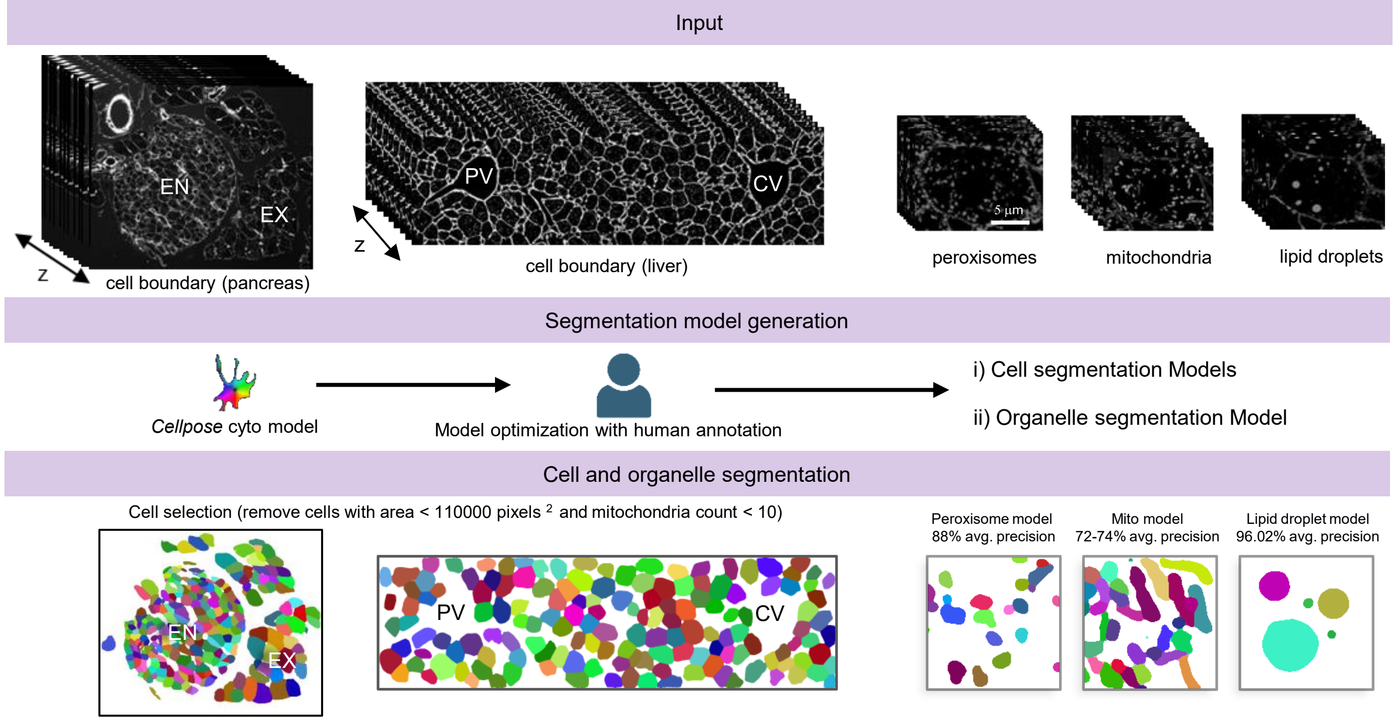


#### C. Feature extraction from segmented masks

Python package for Cellpose segmentation and feature extraction

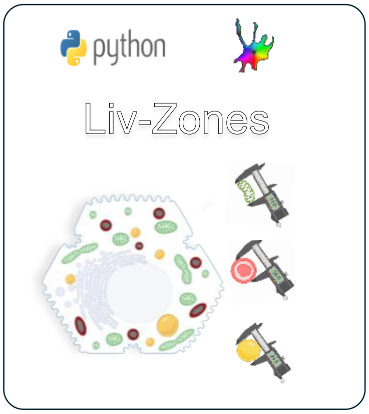


Organelle features extracted

| Relevant Features | Metric used |
|---|---|
| Size | area |
| Shape | aspect ratio |
| Number | density |
| Complexity | 1/solidity |
| Occupancy | Percent total area |
| Distribution within the cell | Distance to the nucleus |
| Acini Position | close to pv = +1<br>Close to cv = -1 |
| Position in tissue | Centroid: x and y-coordinates of center of the mask within the rectangular tissue image |

Visual representation of measured features

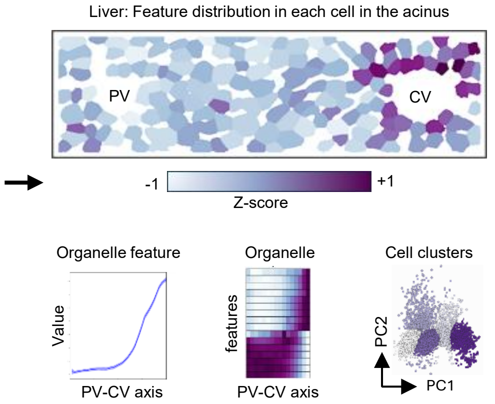

Supplementary Figure 2

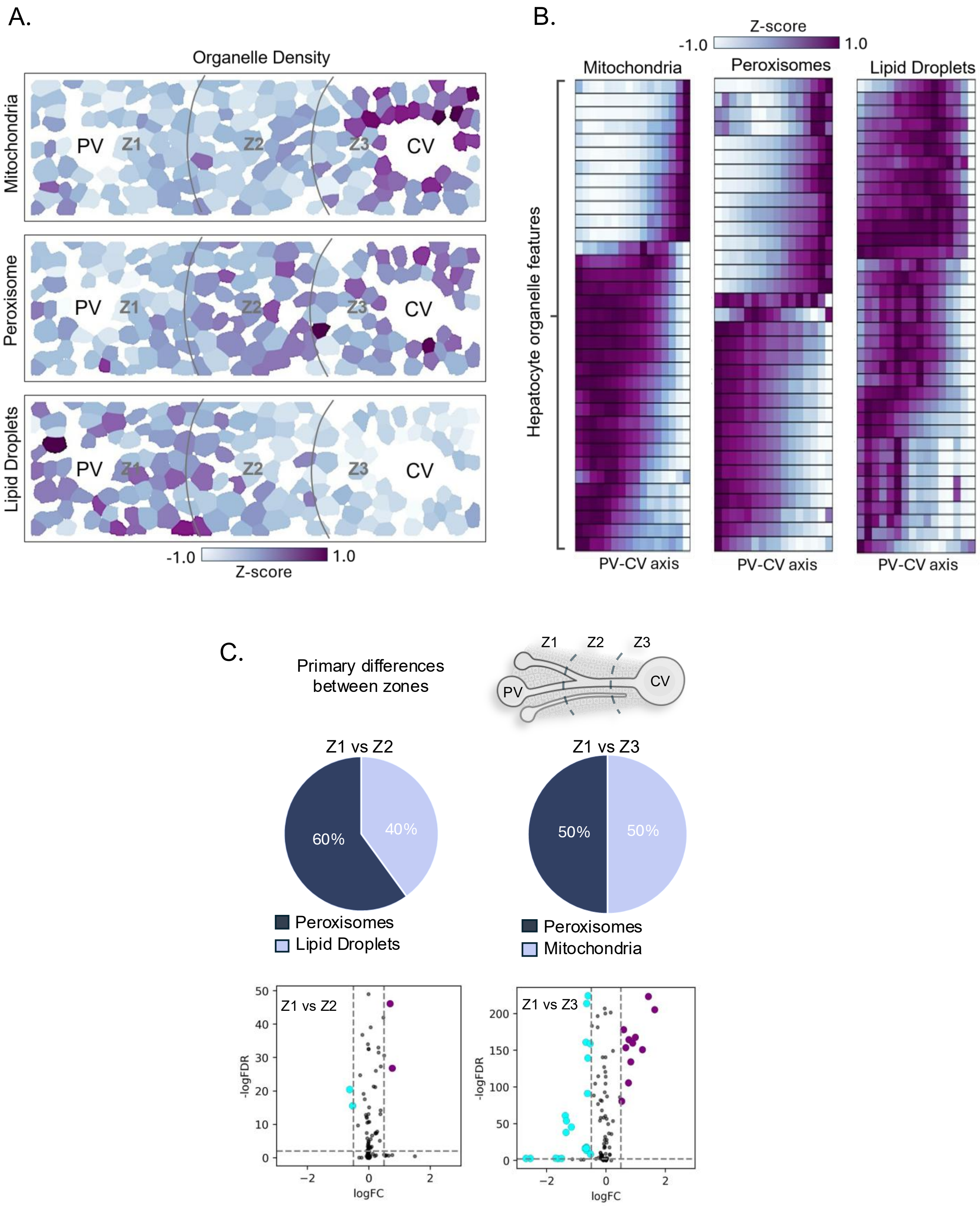

Supplementary Figure 3

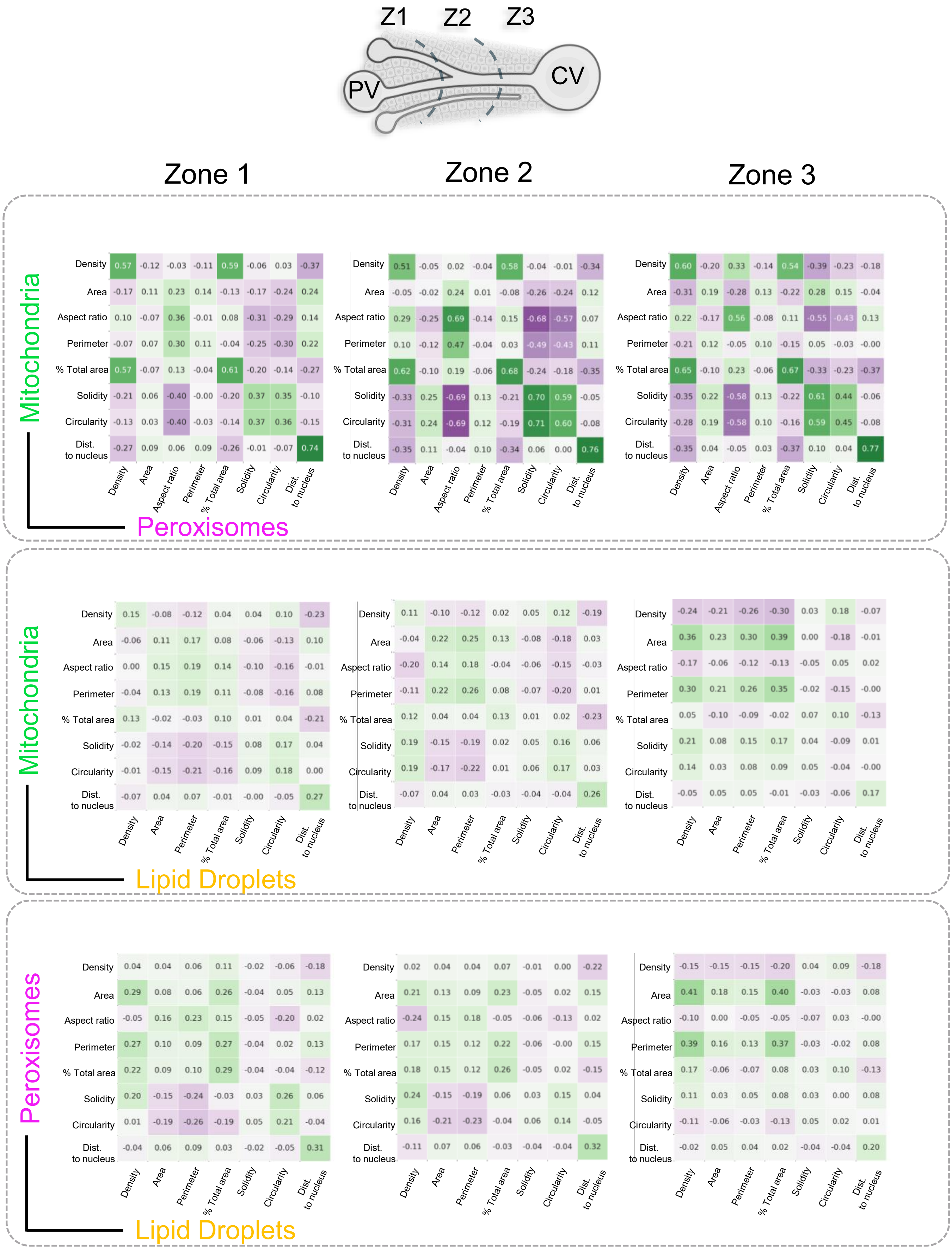

Supplementary Figure 4

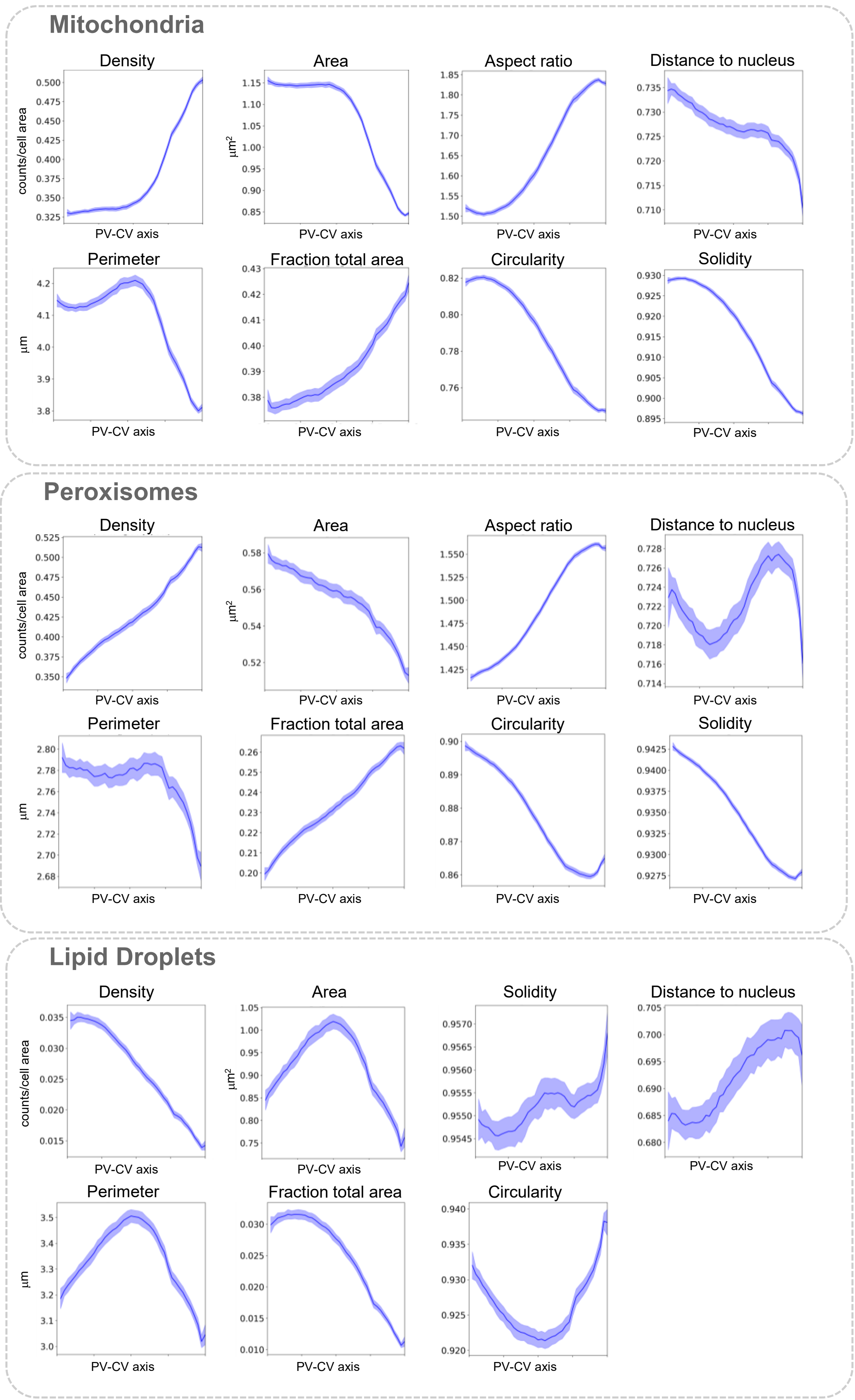

Supplementary Figure 5

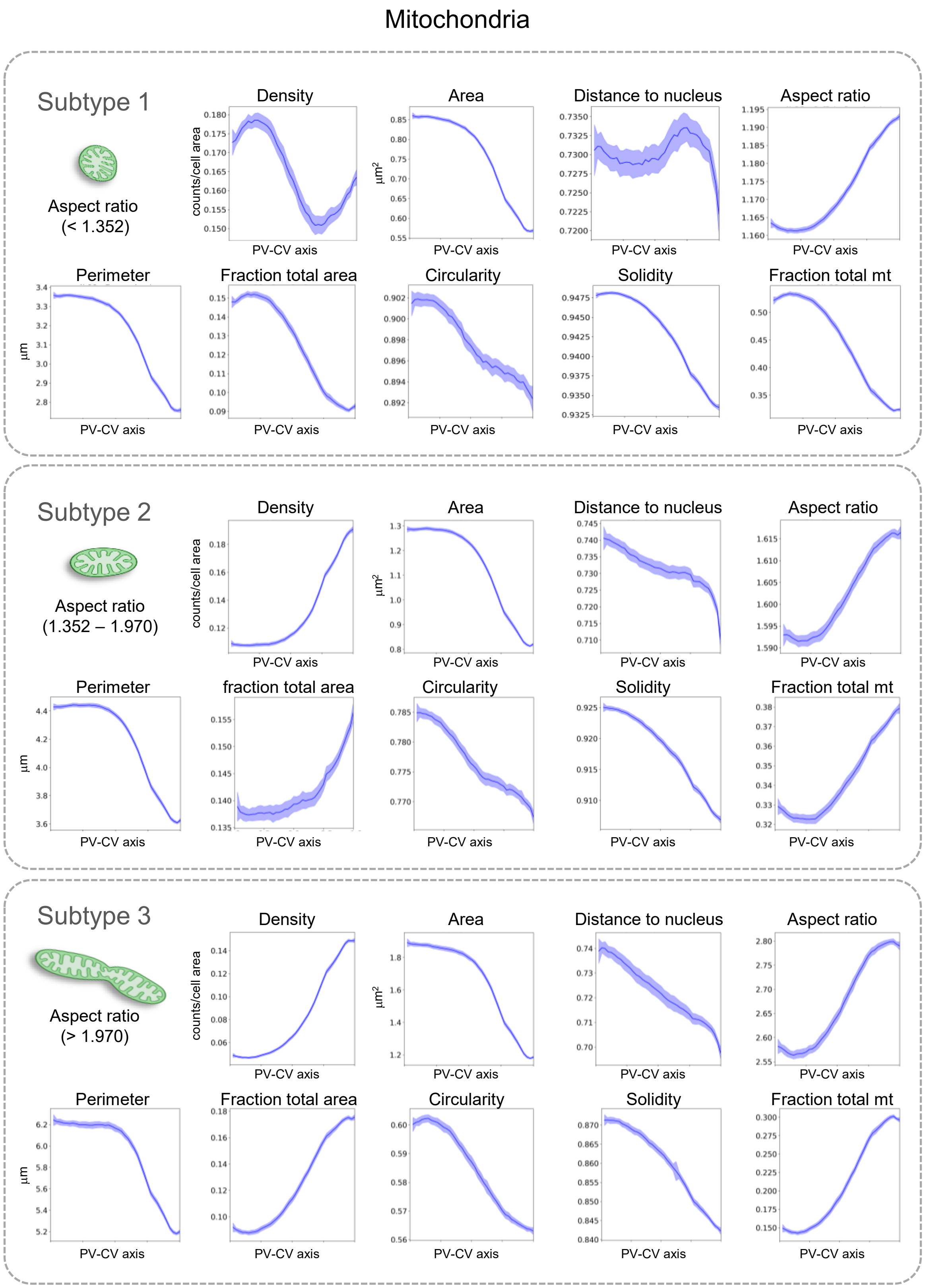

Supplementary Figure 6

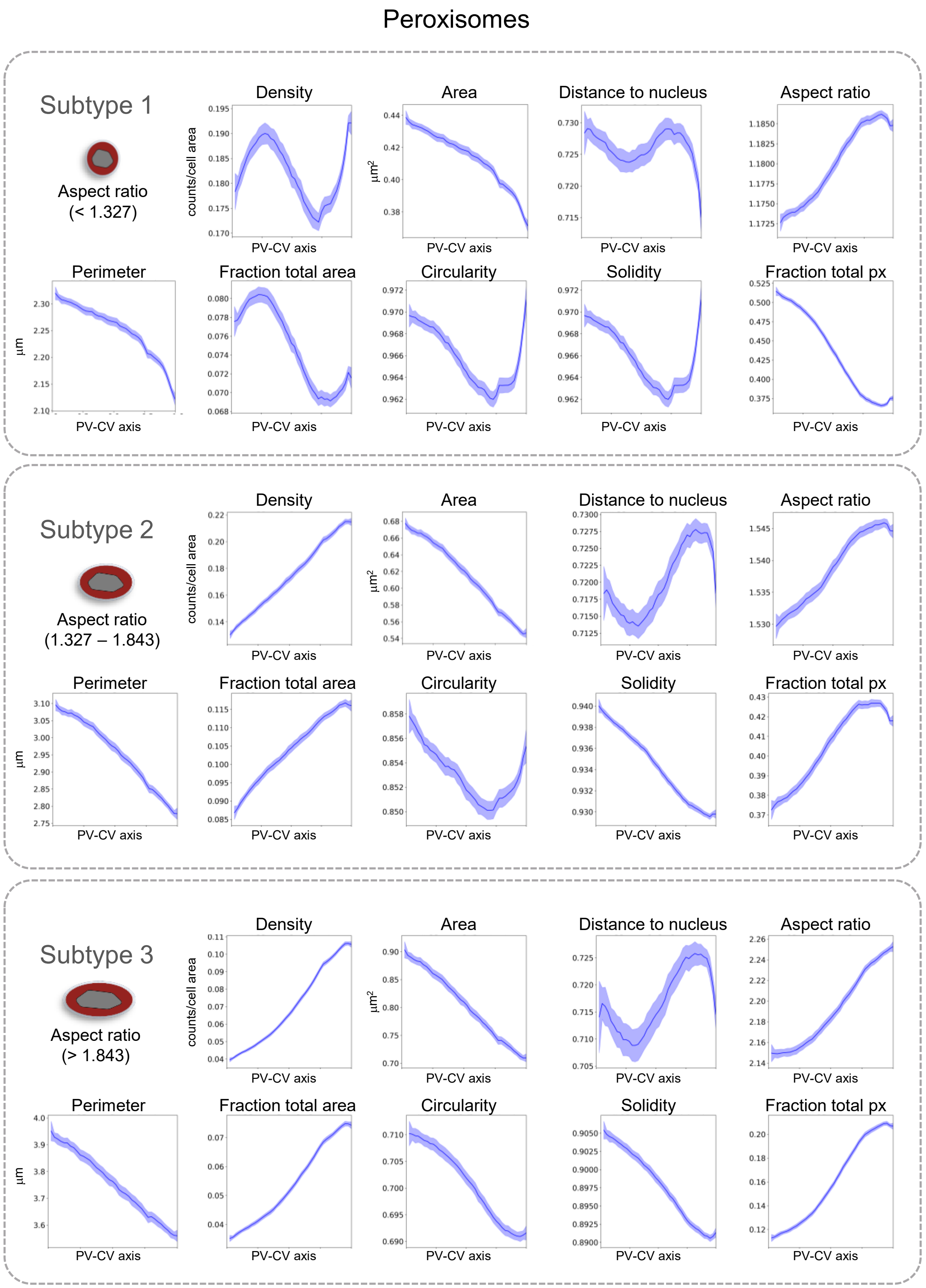

Supplementary Figure 7

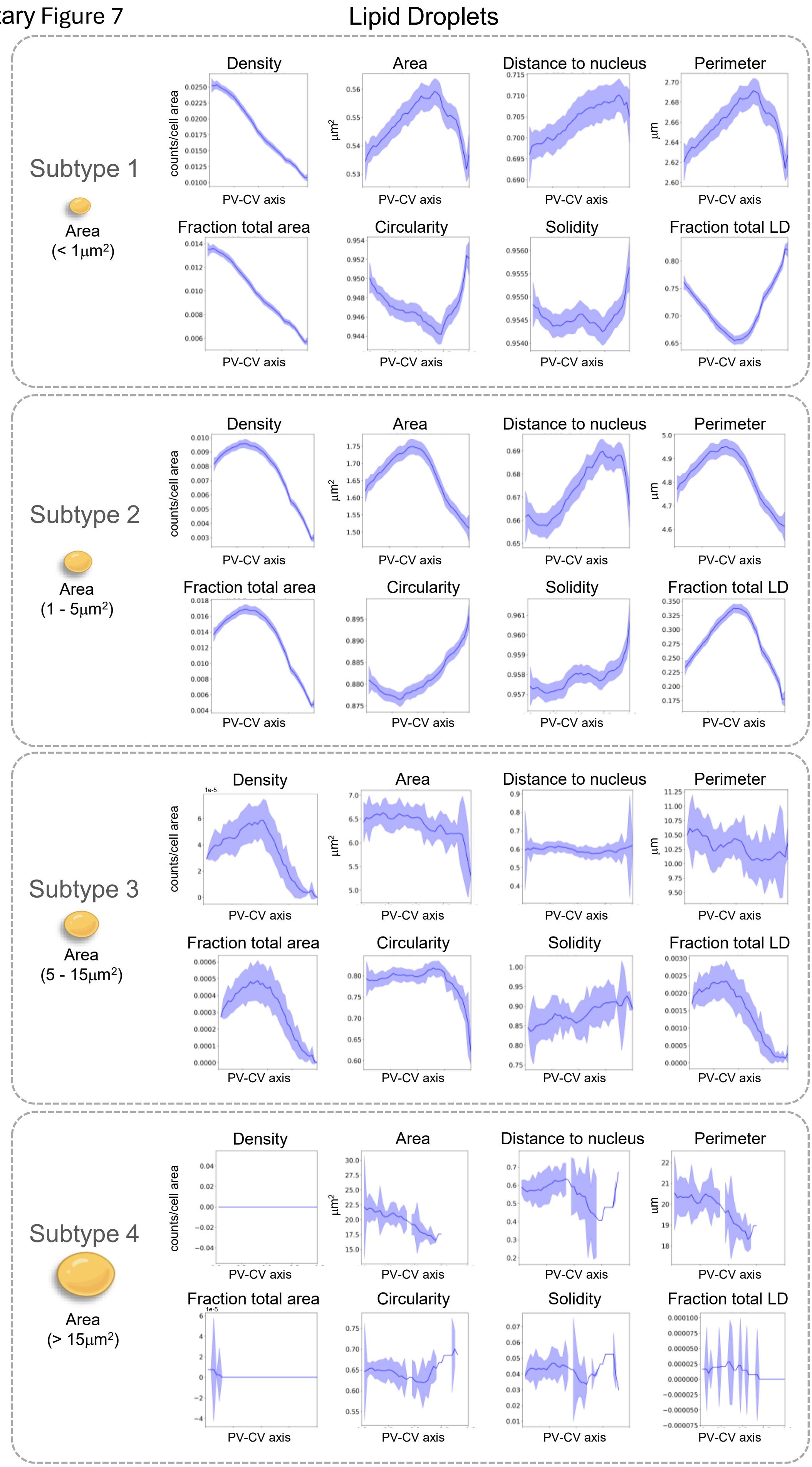

Supplementary Figure 8

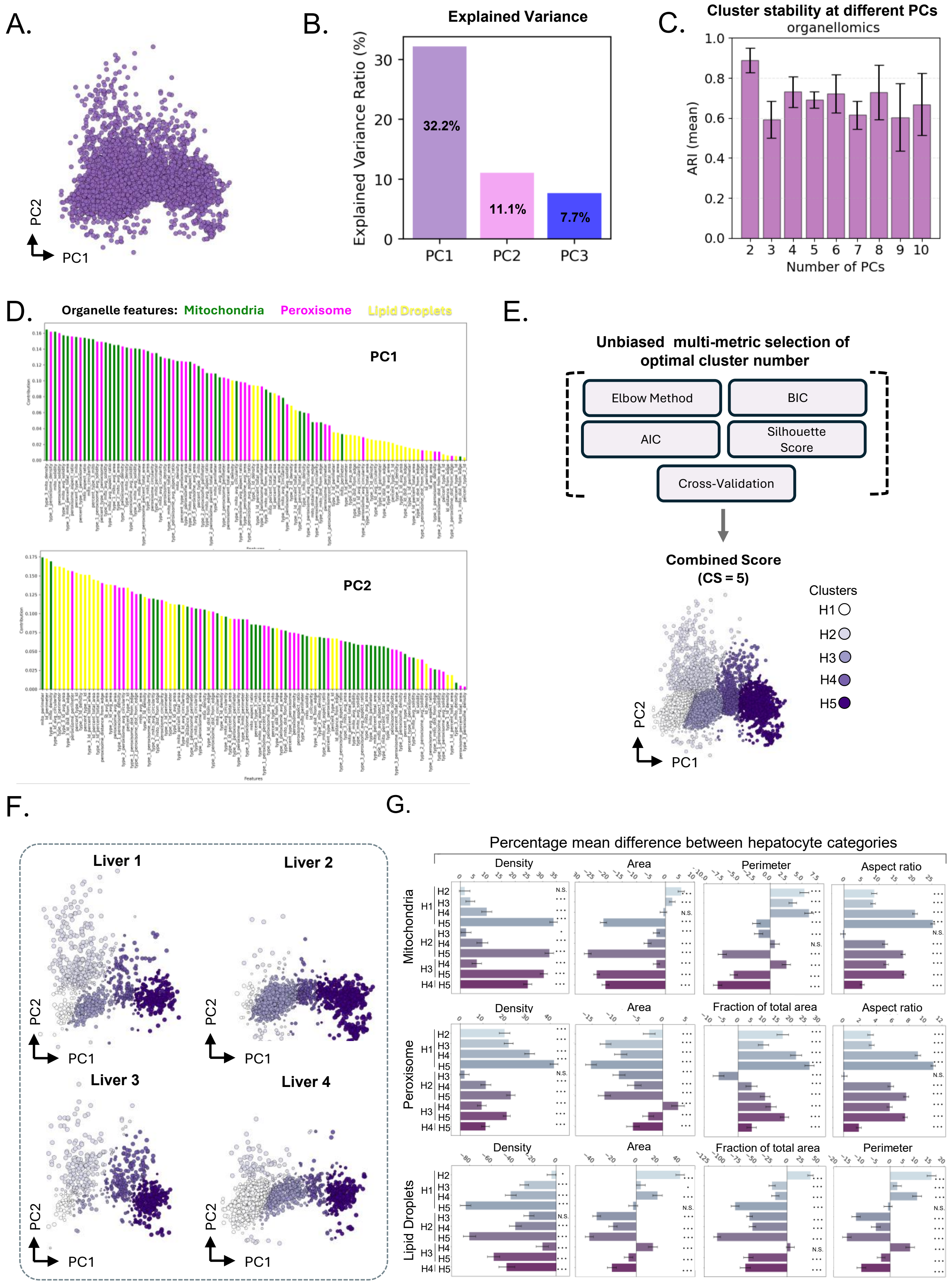

Supplementary Figure 9

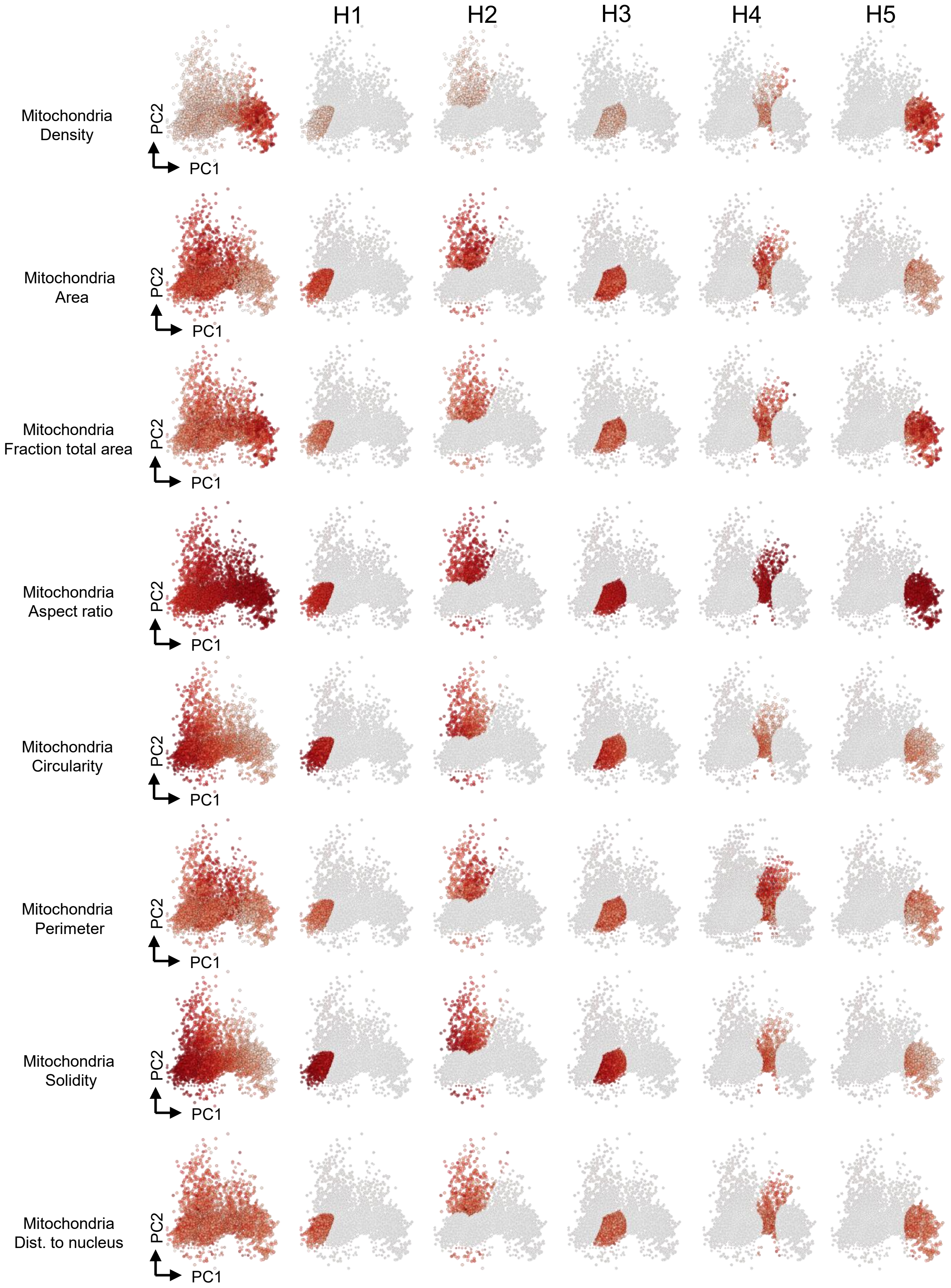

Supplementary Figure 10

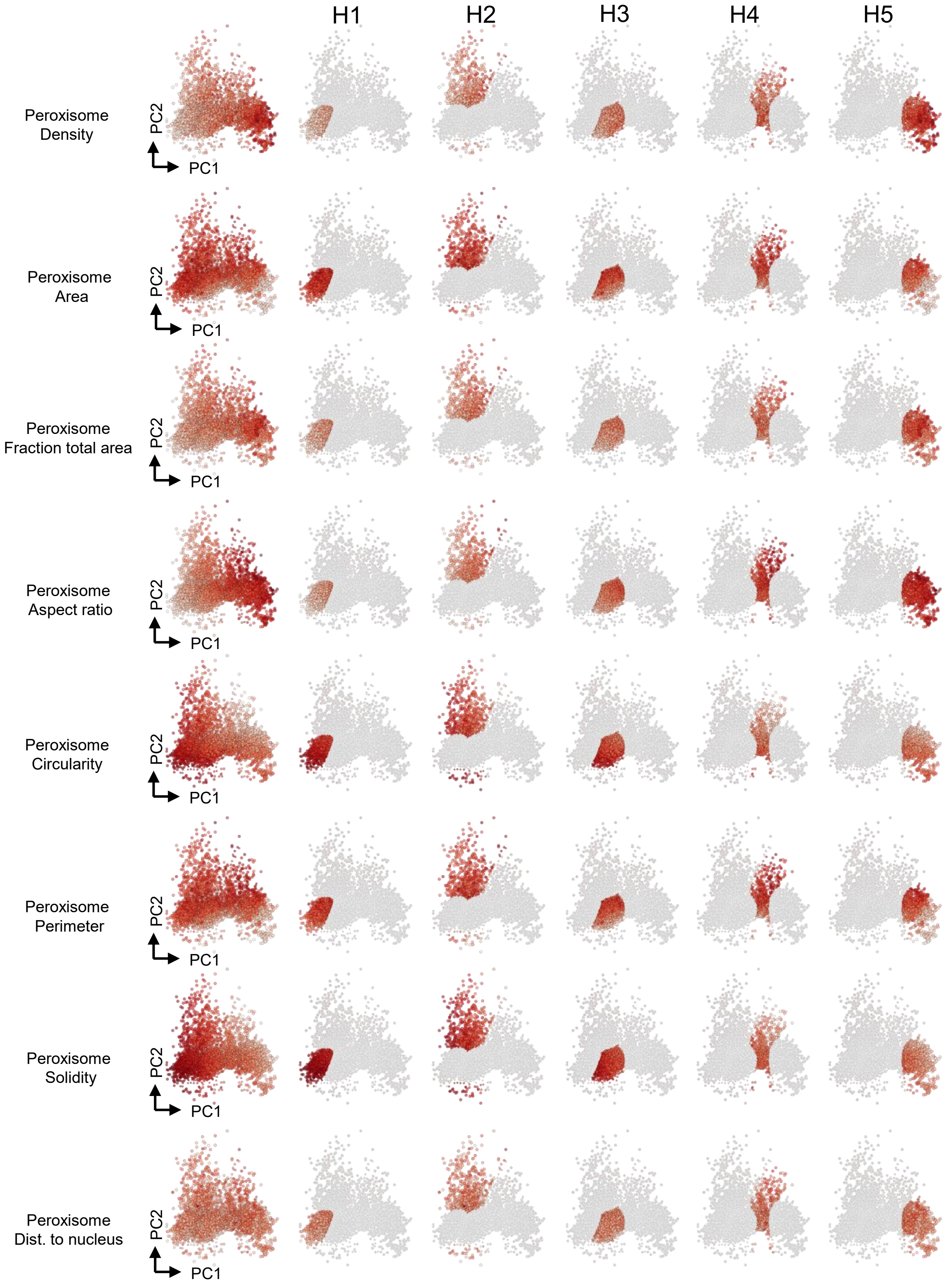

Supplementary Figure 11

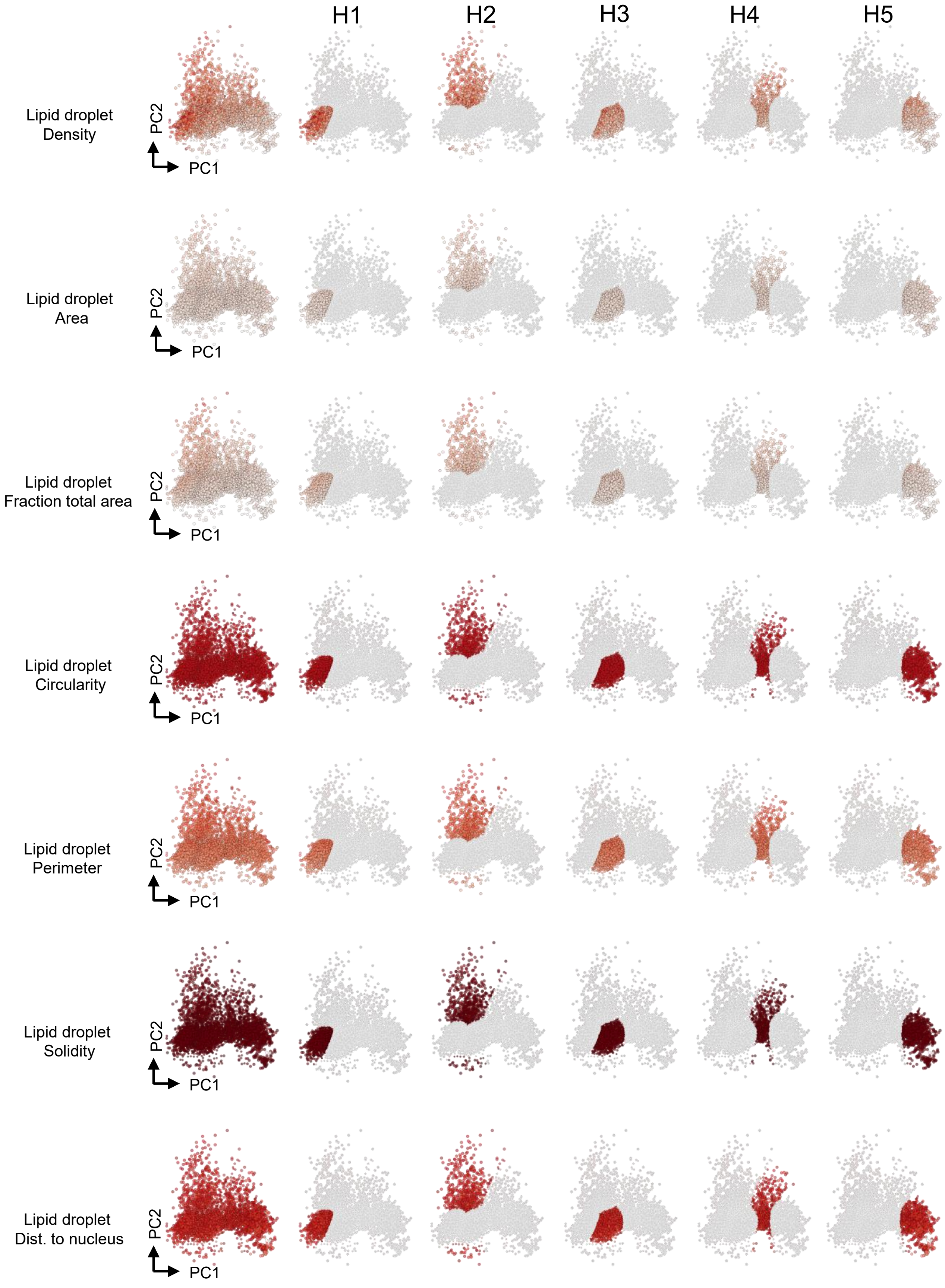

Supplementary Figure 12

A.

| | | Robustness | Separation | Resolution | Effective Resolution | Number of Cells | Number of Features | Number of Clusters | Composite Score | Bootstrap Mean | Bootstrap Std |
|---|---|---|---|---|---|---|---|---|---|---|---|
| Leiden | O | 0.796 | 0.133 | 11.591 | 11.591 | 390 | 98 | 7 | 0.717 | 0.747 | 0.088 |
| | P | 0.698 | 0.115 | 11.06 | 11.06 | 390 | 98 | 7 | 0.43 | 0.501 | 0.08 |
| | T | 0.604 | 0.018 | 14.583 | 14.583 | 390 | 98 | 3 | 0.333 | 0.328 | 0.035 |
| GMM | O | 0.75 | 0.8 | 0.78 | 0.78 | 390 | 98 | 5 | 0.784 | 0.78 | 0.03 |
| | P | 0.75 | 0.78 | 0.77 | 0.77 | 390 | 98 | 2 | 0.781 | 0.78 | 0.02 |
| | T | 0.65 | 0.7 | 0.68 | 0.68 | 390 | 98 | 3 | 0.678 | 0.68 | 0.02 |

B.

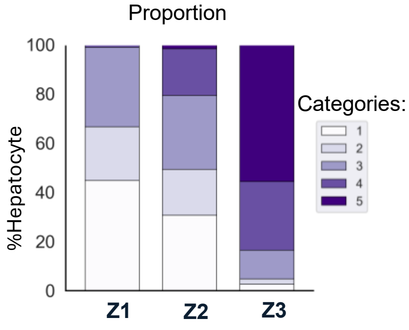


C.

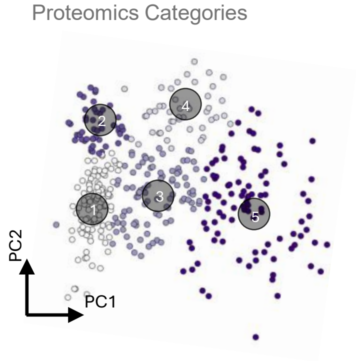


D.

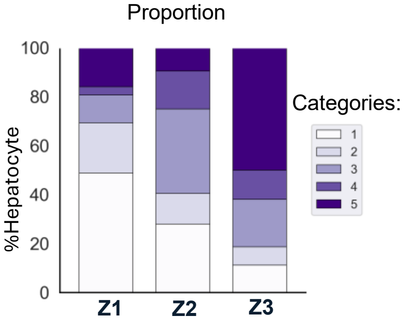

Supplementary Figure 13

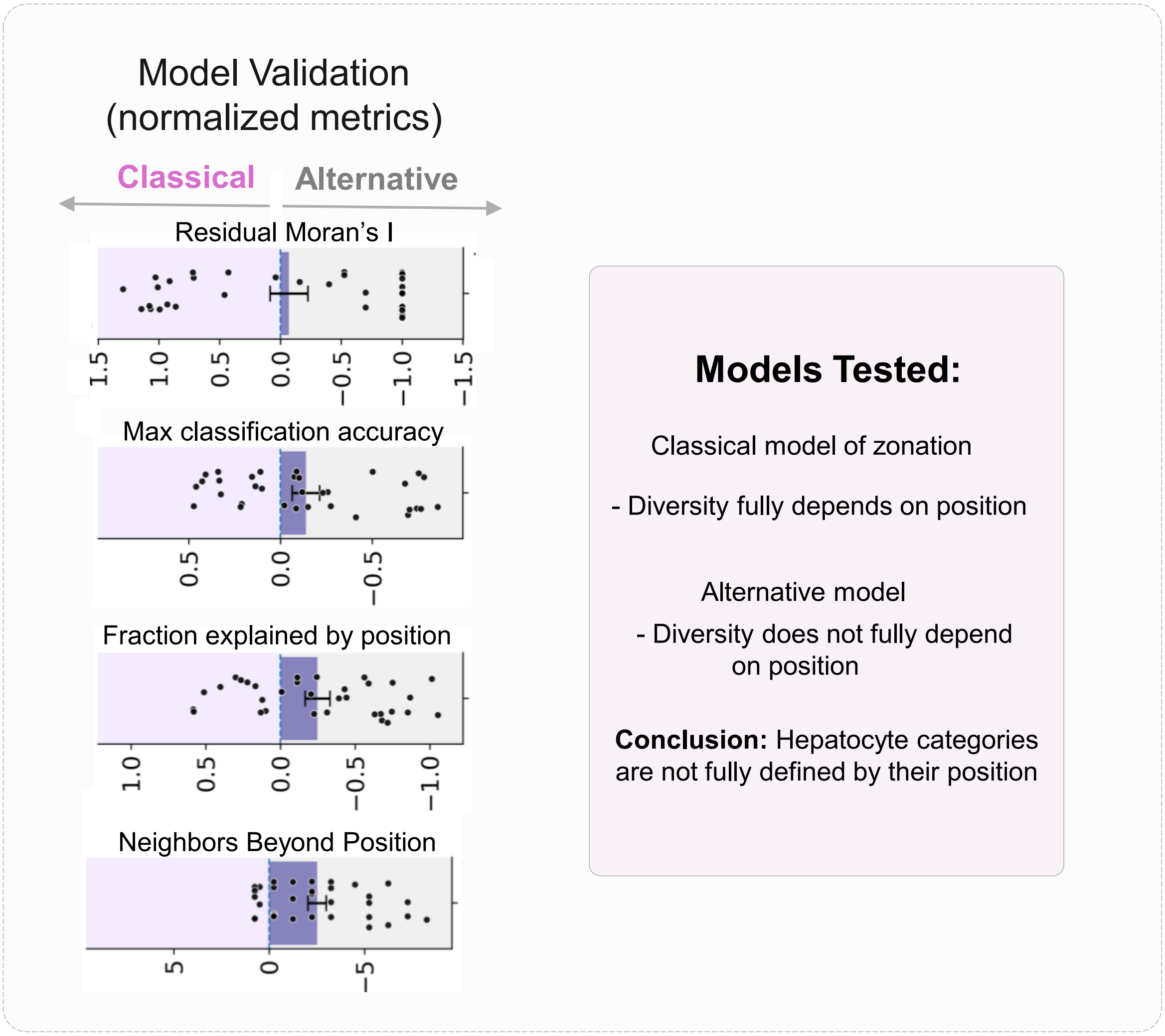

Supplementary Figure 14

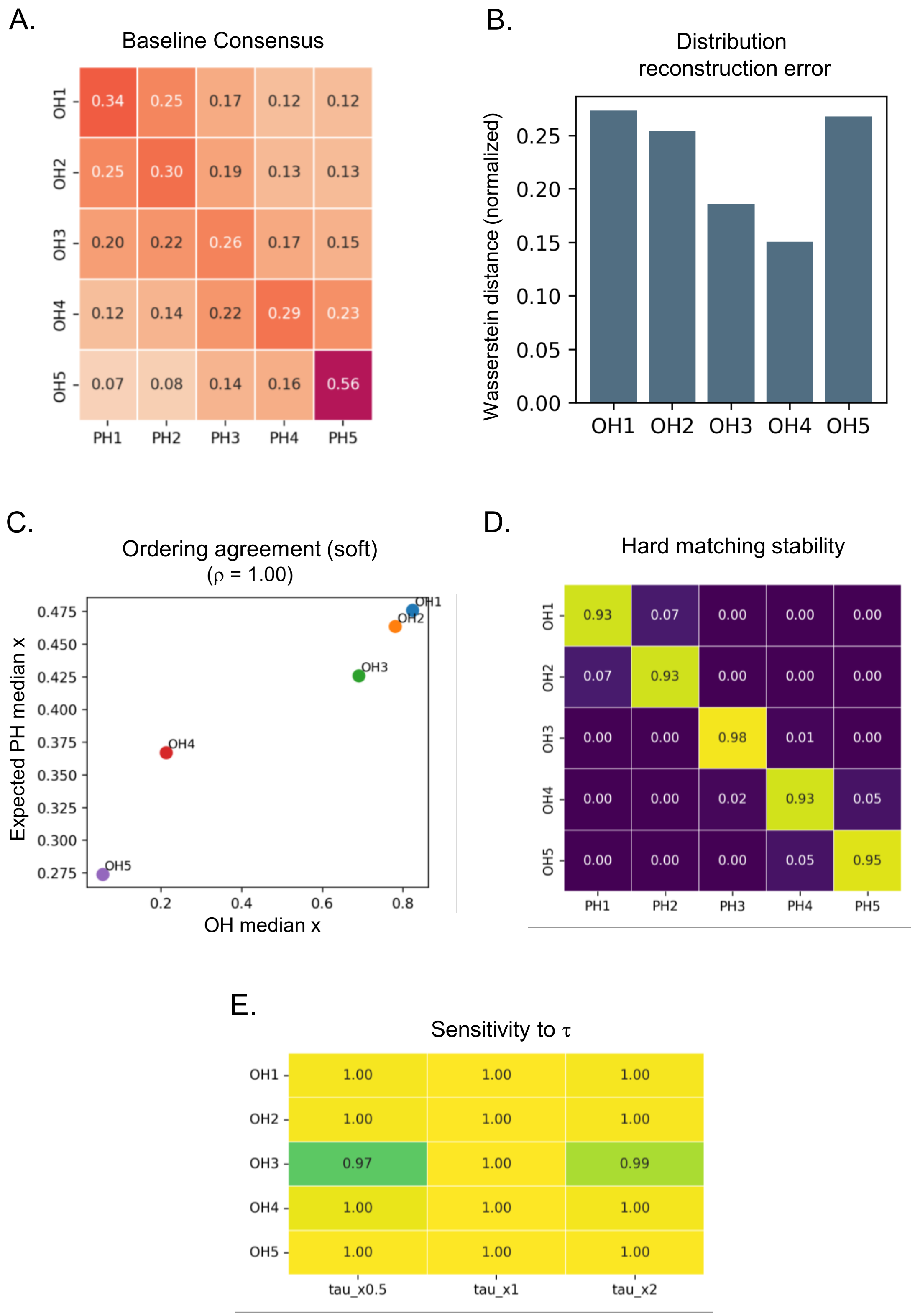

Supplementary Figure 15

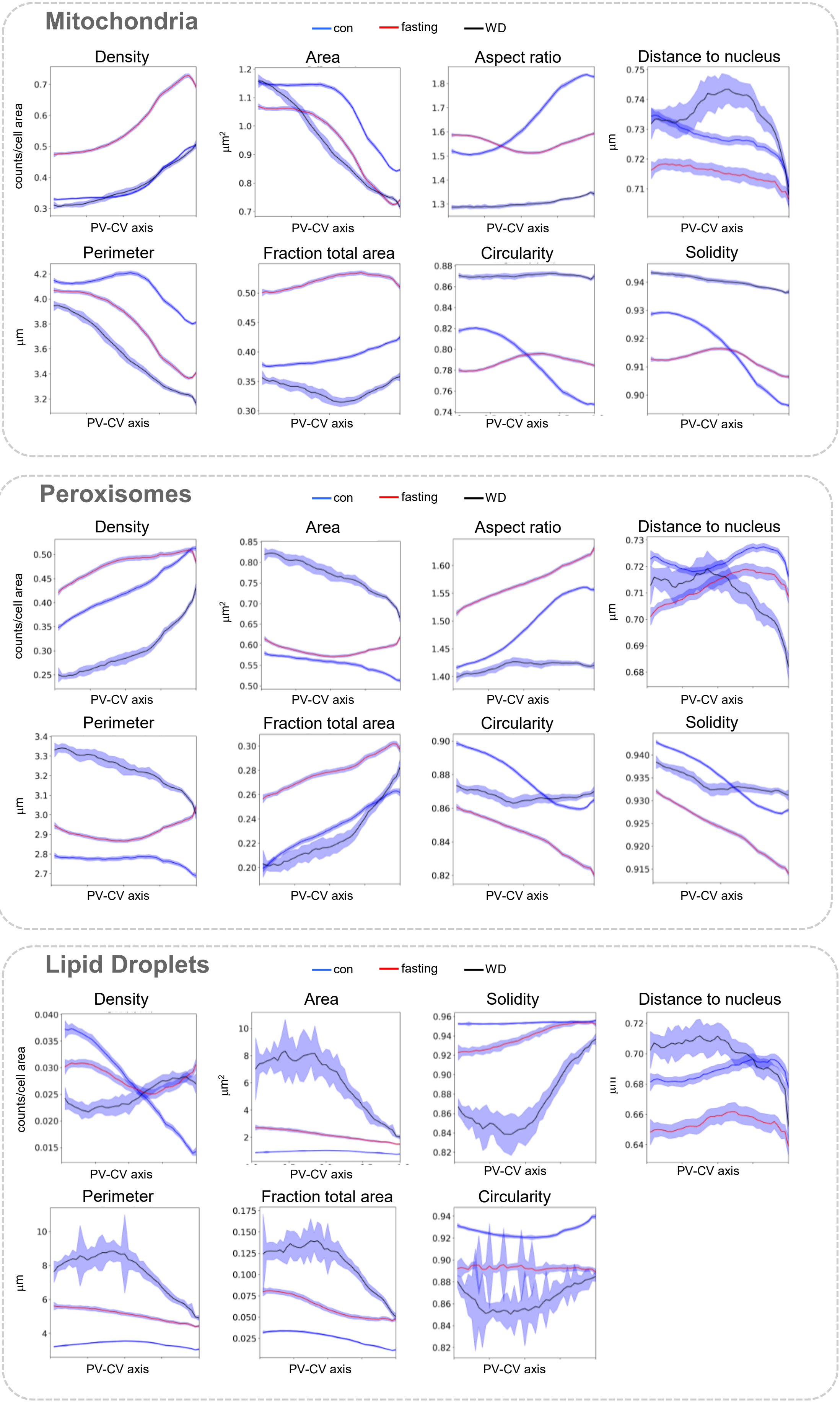

Supplementary Figure 16

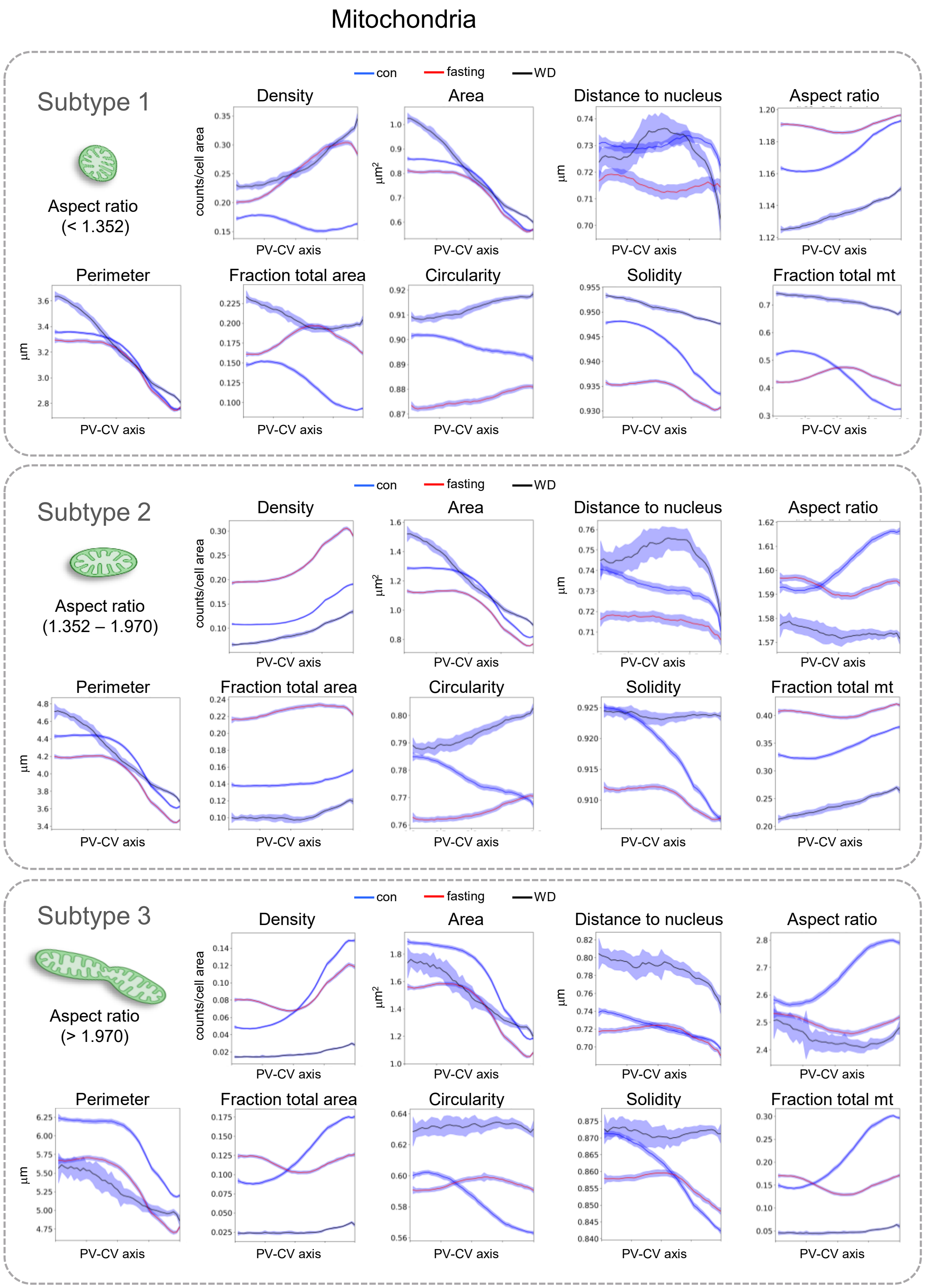

Supplementary Figure 17

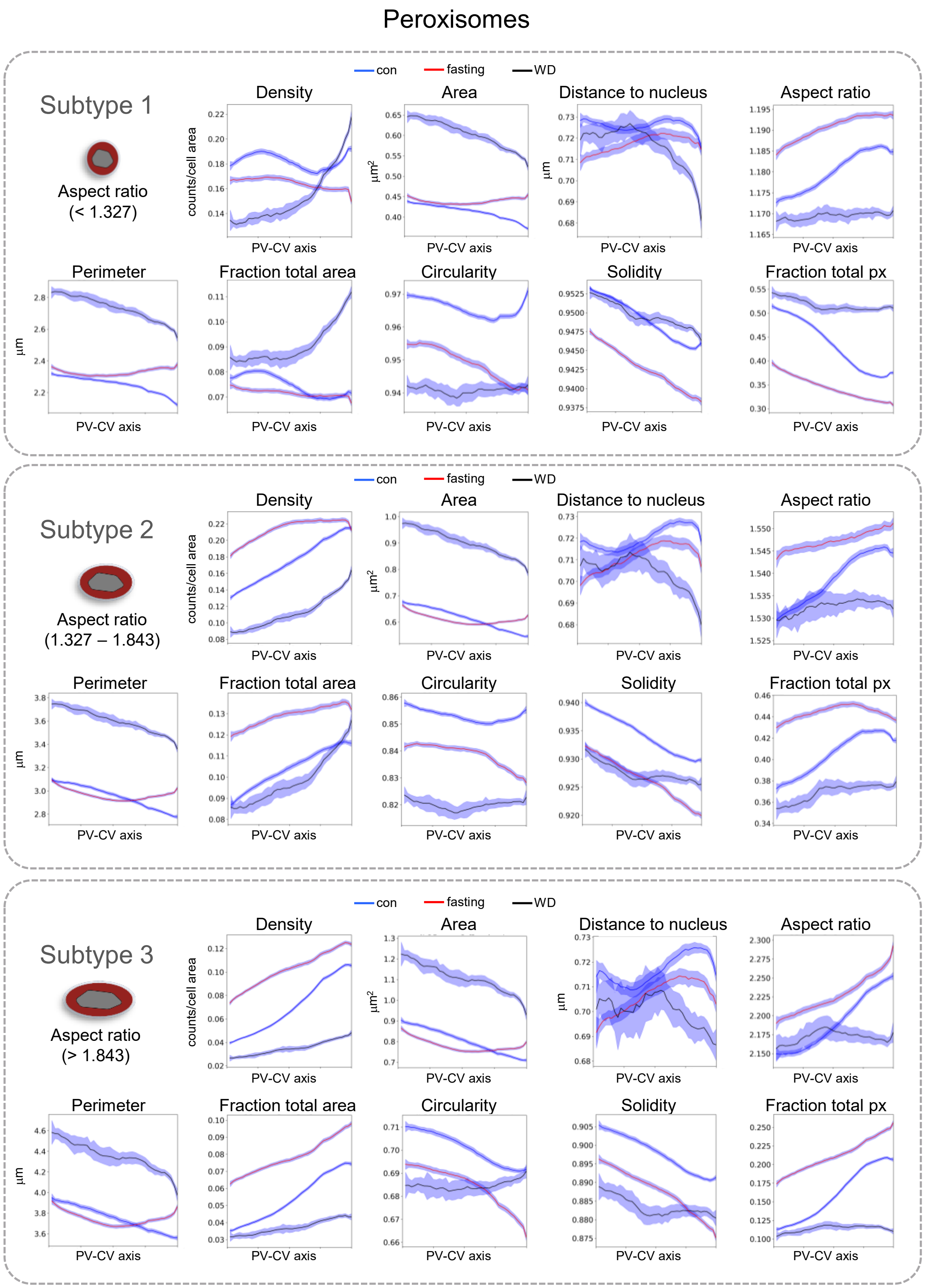

Supplementary Figure 18

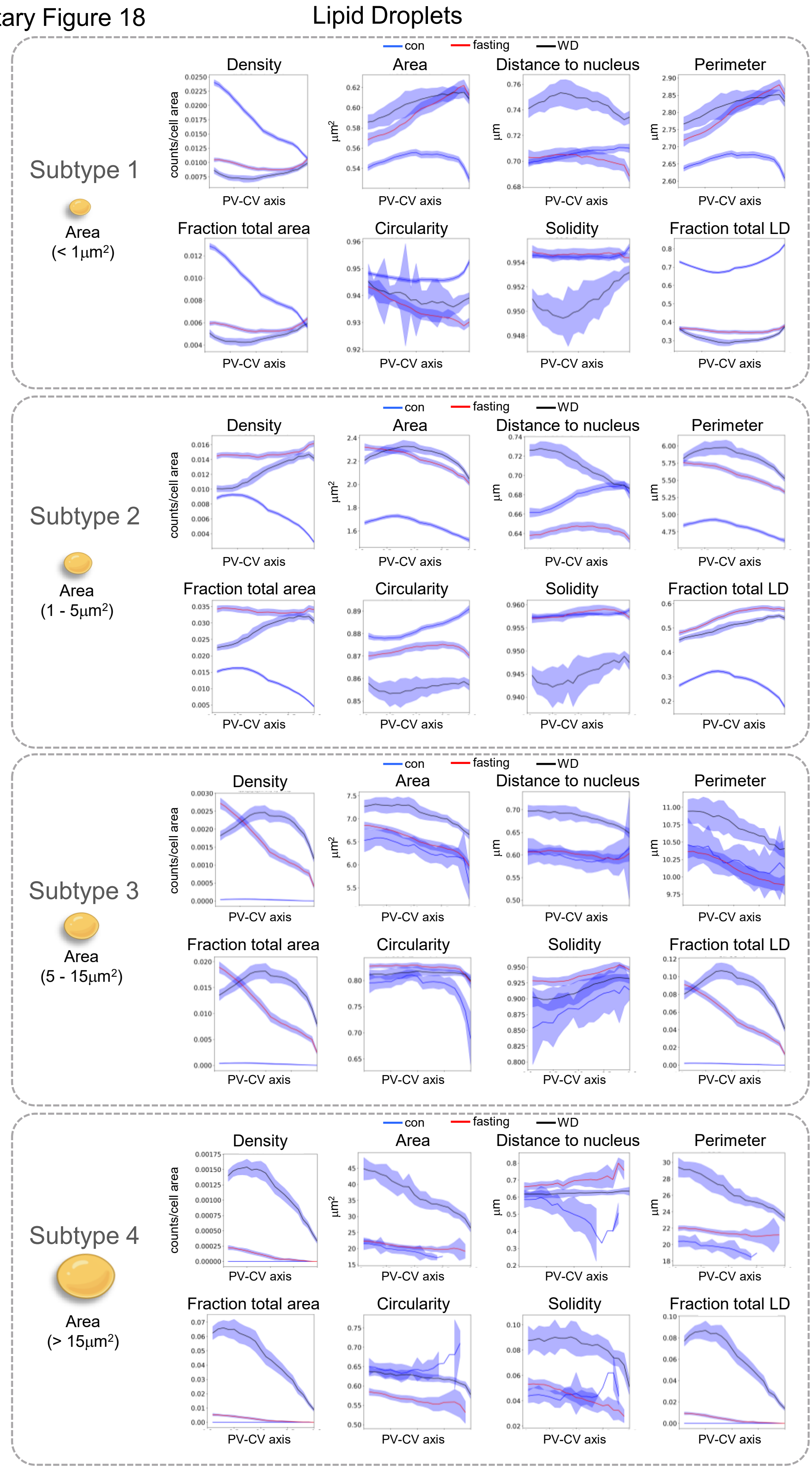

### Supplementary Figure 19

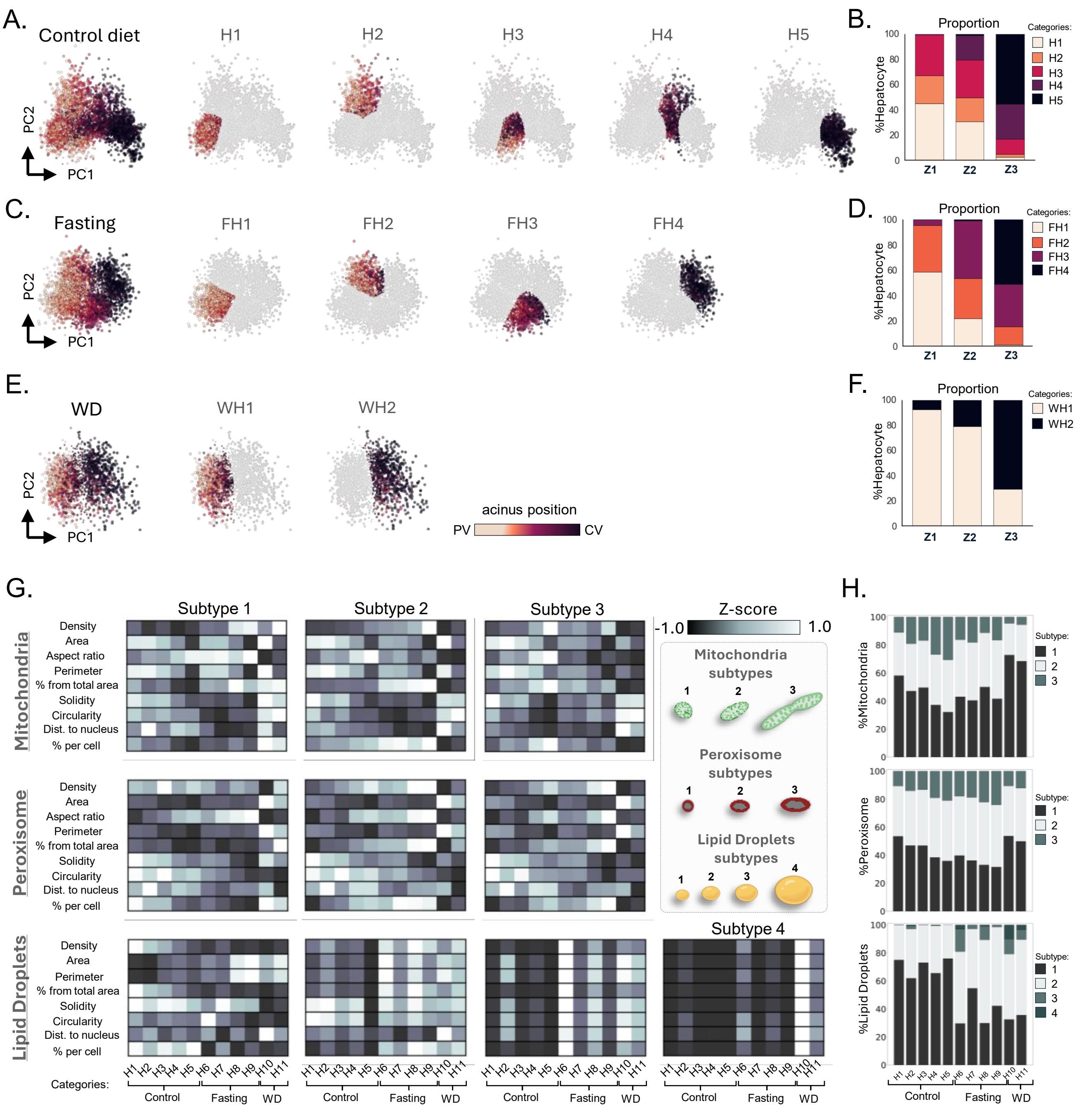

Supplementary Figure 20

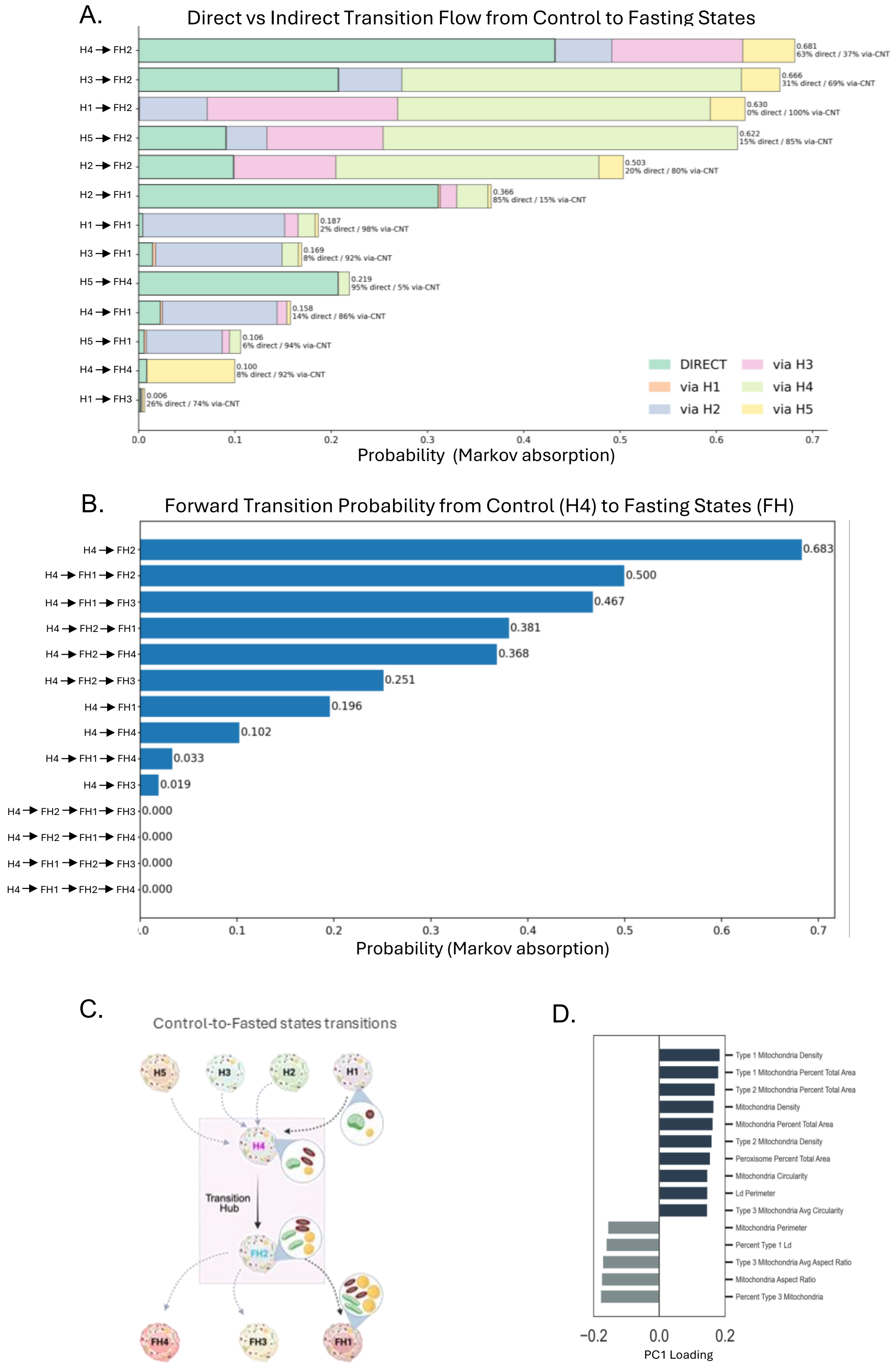

Supplementary Figure 21

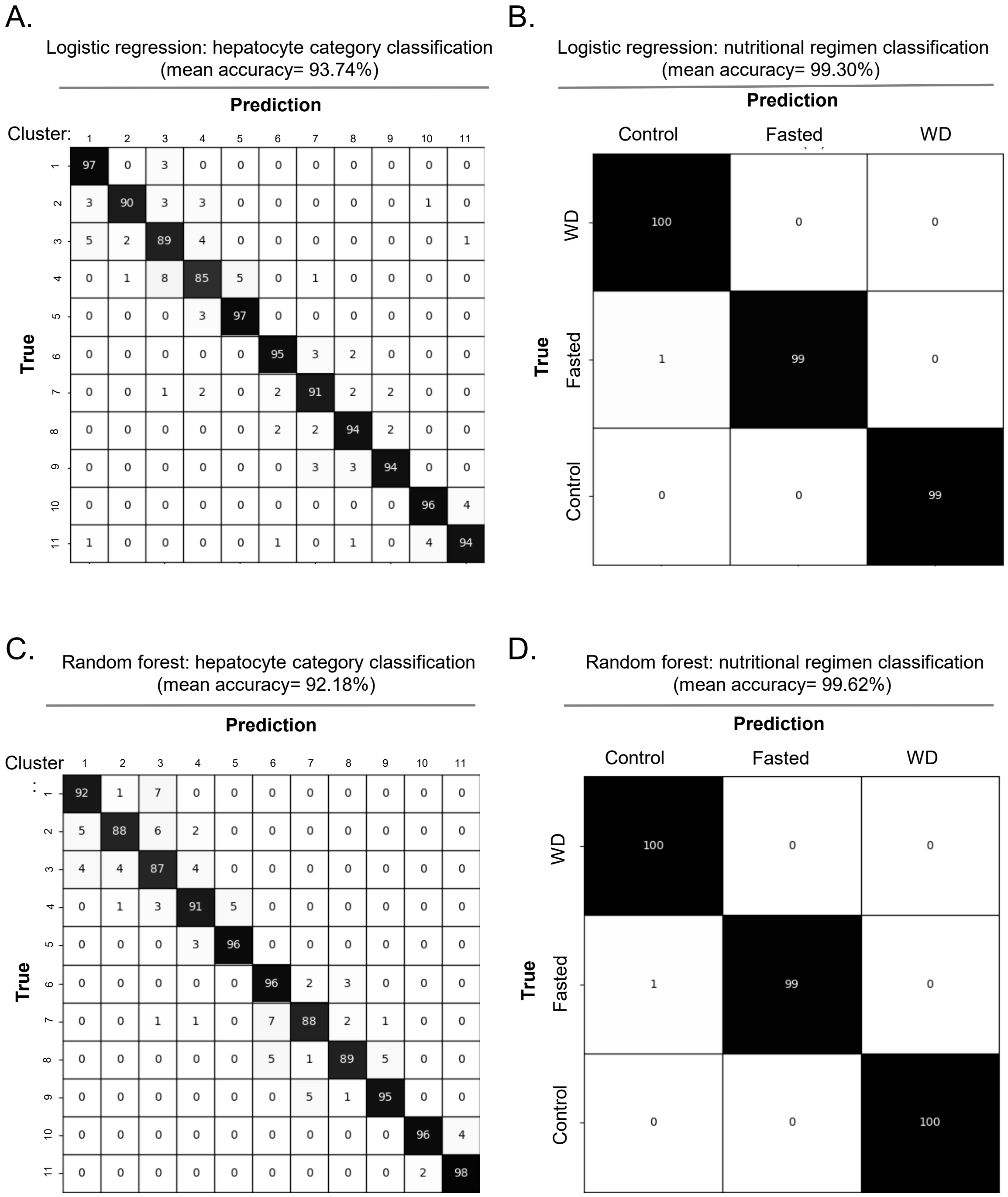
